# Phylodynamics of H5N1 Highly Pathogenic Avian Influenza in Europe, 2005–2010: Potential for Molecular Surveillance of New Outbreaks

**DOI:** 10.1101/020339

**Authors:** Mohammad A. Alkhamis, Brian R. Moore, Andres M. Perez

**Author notes:** Author to whom correspondence should be addressed; (M.A.).

## Abstract

Previous Bayesian phylogeographic studies of H5N1 highly pathogenic avian influenza viruses (HPAIVs) explored the origin and spread of the epidemic from China into Russia, indicating that HPAIV circulated in Russia prior to its detection there in 2005. In this study, we extend this research to explore the evolution and spread of HPAIV within Europe during the 2005–2010 epidemic, using all available sequences of the HA and NA gene regions that were collected in Europe and Russia during the outbreak. We use discrete-trait phylodynamic models within a Bayesian statistical framework to explore the evolution of HPAIV. Our results indicate that the genetic diversity and effective population size of HPAIV peaked between mid-2005 and early 2006, followed by drastic decline in 2007, which coincides with the end of the epidemic in Europe. Our results also suggest that domestic birds were the most likely source of the spread of the virus from Russia into Europe. Additionally, estimates of viral dispersal routes indicate that Russia, Romania, and Germany were key epicenters of these outbreaks. Our study quantifies the dynamics of a major European HPAIV pandemic and substantiates the ability of phylodynamic models to improve molecular surveillance of novel AIVs.

## 1. Introduction

The epidemiology of the H5N1 highly pathogenic avian influenza viruses (HPAIVs) is characterized both by the far-reaching economic impact for commercial poultry production, and also by the risk of spread into mammalian species, including humans. Transmission into humans has mainly been attributed to direct contact of susceptible individuals with infected poultry, including activities associated with the slaughter and dressing of poultry for human consumption [1]. The maintenance and spread of avian influenza (AI) infection in domesticated poultry are mainly associated with the establishment of the virus in free-range (or backyard) poultry and its subsequent introduction to commercial animals via live-bird markets. Furthermore, wild birds are the natural reservoirs of the virus, and the movement of wild birds infected with HPAIV is known to play an important role in the spread, circulation, and maintenance of the virus through either direct or indirect contact with poultry [2–4].

The ancestral strain of H5N1 HPAIV, referred to as A/goose/Guangdong/1/96, emerged in commercial domesticated geese in the Guangdong province of China in 1996 [5]. Since then, the strain has undergone numerous genetic changes that have given rise to several distinct lineages, also referred to as clades [6]. To date, ten H5N1 HPAIV clades have been identified based on comparisons of 859 hemagglutinin (HA) gene sequences [7]. These viruses are now widespread in countries throughout Asia, Europe, and Africa. Between 2004 and 2005, a new H5N1 HPAIV strain (sub-clade 2.2) emerged that caused severe outbreaks in wild birds and poultry in northwestern China [6, 8]. In Europe, HPAI H5N1 was first detected at the end of 2005 in both poultry (in Russia, Romania, and Turkey) and wild birds (in Croatia) [2, 9–11]. In January 2006, the first human infection outside Southeast Asia was reported in Turkey. In February 2006, the first of occurrence of HPAI H5N1 in Northern Europe was identified from a wild mute swan (*Cygnus olor*) in Germany in the southwestern part of the Baltic Sea [11,12]. Between 2006 and 2010, the epidemic spread through 13 European Union (EU) member states located eastward of a line extending from south-eastern Sweden to south-western Italy. Both poultry and wild-bird populations were infected in Austria, Bulgaria, Czech Republic, Denmark, France, Greece, Hungary, Poland, Slovak Republic, Slovenia, Switzerland, and the UK [10, 13]. Phylogenetic analysis of HPAIVs isolated from European outbreaks distinguished at least two different sub-lineages of the 2005 Qinghai strain, which confirmed that the virus spread from southeastern Asia into Europe [10–13].

Since 2005, surveillance of avian influenza viruses in poultry and wild birds has become compulsory in the EU. Member states have conducted extensive molecular surveillance for AI, with most of the studies focused on genetic analysis of partially or fully sequenced AI viral genomes [14–18]. Molecular epidemiological studies of H5N1 HPAIV sequence data isolated between 2005–2010 in EU member states, for example, were focused on the molecular characterization, genetic classification, identification of genetic shifts and drifts, and the degree of relatedness to previously isolated viruses. Phylogenies of H5N1 HPAIV isolated in the EU were estimated using traditional parsimony, neighbor-joining, and maximum-likelihood methods. However, those methods typically ignore various sources of uncertainty, including uncertainty associated with estimates of the phylogenetic relationships, divergence times, and history of geographic dispersal [19]. Furthermore, epidemiological studies that explored risk factors and the spatial and temporal evolution of H5N1 HPAIV in the EU between 2005– 2010 were mainly conducted separately from phylogenetic studies [20–24]. Phylodynamics is an emerging field that aims to characterize the joint evolutionary and epidemical behavior of rapidly evolving infectious diseases using tools borrowed from the field of phylogenetics [25]. The probabilistic models used in the phylodynamic approach are mostly pursued in a Bayesian statistical framework [26]. This approach treats parameters of the phylodynamic model as random variables, such that each parameter is described by a specified prior probability distribution (and a corresponding inferred posterior probability distribution). Accordingly, the Bayesian approach provides a natural way to estimate (and accommodate) uncertainty in the phylodynamic model parameters, including the virus phylogeny, divergence times, and history of geographic spread [27].

Recent evidence suggests that Bayesian methods may be more accurate in assessing HPAIV evolution than traditional phylogenetic methods. For example, Lemey *et al.* (2009) [27] adopted a Bayesian phylodymanic approach to re-analyze a H5N1 HPAIV dataset initially studied using conventional methods [28, 29]. Results of the Bayesian analysis did not support the earlier conclusion regarding the epidemiological link between Guangdong and Indonesia, but instead suggested that the ancestral strains of the Indonesian outbreaks originated from the Hunan province in south central China [27]. Furthermore, these studies demonstrated the potential of Bayesian phylogeographic methods to estimate the posterior probability of each geographic location at any point in the phylogenetic tree, and the ability for phylogenetic parameters to be estimated from different genomic segments of the H5N1 HPAIV genome without assuming that the sequences shared an identical phylogenetic history [27].

Here, we extend the landmark study of Lemey *et al.* (2009) to estimate the history of HPAIV dispersal into and within Europe after its initial detection in Russia in 2005. Our data comprise 277 H5N1 HPAIV hemagglutinin (HA) and neuraminidase (NA) gene sequences (referred to as isolates) collected between 2005 and 2010 in Europe and Russia. We adopt a discrete-trait phylodynamic model to estimate both the history of viral migration between geographic areas, and also to infer the movement of the virus among host species (wild versus domesticated birds). We also estimate temporal changes in viral population size, and discuss the potential of phylodynamic methods to improve AI surveillance. Our study provides quantitative estimates of the mechanisms that led to the rapid spread of HPAIV throughout Europe in 2005–2010 and further illustrates the potential of this approach to improve the prevention and control of novel emerging AIVs.

## 2. Results and Discussion

Our analyses of the viral population dynamics for both HA and NA genes reveal a strong increase in genetic diversity (and, therefore, effective population size) in mid-2005 followed by a second increase in early 2006 (Figure 1B and C). This coincides with the highest frequency of reported cases of H5N1 infections during the epidemic, when several independent introductions of H5N1 viruses belonging to sub-clade 2.2 have been confirmed (Figure 1A)[15, 16, 30, 31]. The relatively high viral genetic diversity during the early stage of the epidemic is consistent with the introduction of the virus into a novel region, with a naïve population, where it was exposed to selection pressure imposed by environmental and demographic factors [10, 23, 31]. We also inferred a third (and relatively minor) change in viral genetic diversity during mid-2007, which coincides with the entry of the virus into new EU countries. From 2005 to 2007 conditions seem to have been relatively favorable for the virus to perpetuate in the susceptible population and, given the genetic diversity, reassortment may have occurred during this time frame. The viral genetic diversity is then inferred to decline toward 2007 and to stabilize until 2010. The inferred decrease of viral genetic diversity coincides with the decline of the epidemic, suggesting that prevailing epidemiological factors, such as host density or weather conditions, in 2007–2010 were not sufficient to favor the establishment of the newly emerging H5N1 strain in the population. In contrast to typical AIV epidemics (which exhibit seasonal variation in viral genetic diversity) the genetic diversity of the H5N1 virus did not exhibit seasonal variation in genetic diversity during the 2005–2010 period of the epidemic in Europe and Russia, which suggests that transmission of the virus was influenced by factors different from those typically observed in AIV epidemics.

**Figure 1.**
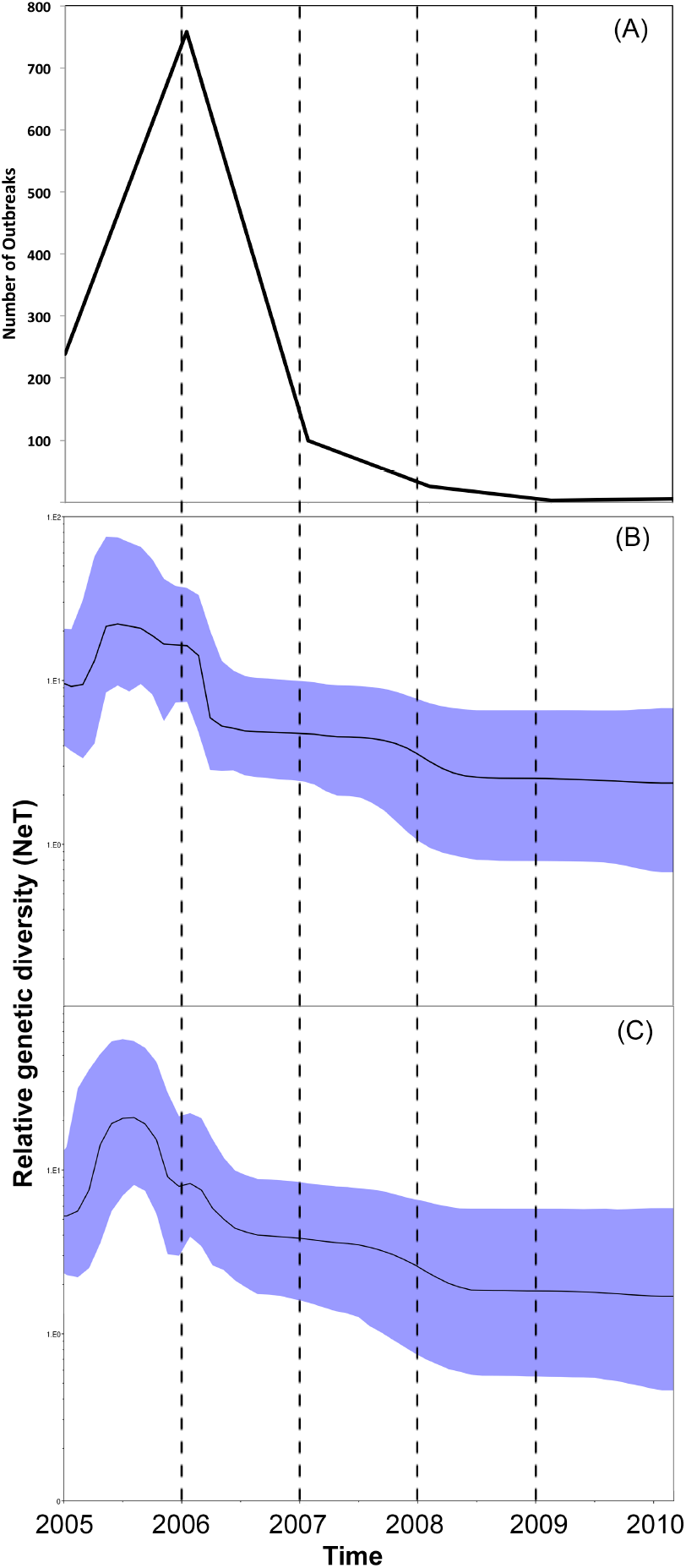
**(A)** Temporal distribution of H5N1 HPAI outbreaks (per year) in domesticated poultry and wild birds in Europe and Russia from between 2005 and 2010. **(B)** and **(C)** Bayesian Skyline plots (BSP) illustrating temporal changes in the relative genetic diversity of H5N1 HPAIV isolates from outbreaks in Europe and the Russia between May, 2005 and June, 2010 estimated from the HA and NA gene sequences. Line plots summarize estimates of the effective population size (*N*_*e*_*T*), a measure of genetic diversity, for HA (above) and NA (below) gene regions; the shaded regions correspond to the 95% HPD.

Our analyses indicate that domestic birds were the most probable ancestral host type for both genes. Domestic birds are inferred to be the most probable host type along many branches of the HA and NA phylogenies, suggesting that the evolution of the H5N1 HPAIV virus largely occurred in association with domestic-bird populations. By contrast, the large proportion of wild-bird isolates (Figure 2) and the inferred high transition rates between host types (Table 1) suggest that wild birds also played an important role in viral dispersal between host populations and geographic areas. This finding is consistent with a number of alternative hypotheses, including scenarios where (1) domestic birds were the source of the spread of H5N1 HPAIV into Europe, or (2) where the HPAI strains were first transmitted from non-EU wild birds to non-EU domestic birds (with little or no genetic change), and later transmitted from non-EU domestic birds to EU domestic birds (Figure 2). The first scenario may be explained by the illegal introduction of domestic birds into Europe, which has been implicated in the spread of other diseases within some European countries [34]. The second scenario assumes very low selective pressure on the virus within—and rapid transmission of the virus among—wild-bird populations, which would explain the high mortality in wild-bird populations and the reported prevalence of the virus along migratory routes [33]. These two seemingly contradictory scenarios may be reconciled as follows: H5N1 HPAIV was initially transmitted from non-EU wild birds to domestic poultry as a less virulent and non-reassortant strain of the virus. Subsequently, the virulent strain of H5N1 HPAIV that caused the EU epidemic emerged as a reassortant strain from poultry, as suggested elsewhere [32]. The high density of poultry populations may have resulted in elevated contact and transmission rates, ultimately enhancing opportunities for viral evolution in poultry compared to wild birds. Additionally, the three sequences isolated from non-avian species were obtained from hosts that prey on birds (two from cats and one from a marten), which is both consistent with a wild-bird origin of infection (Table 1) and with the results of previous studies [11, 35].

**Figure 2.**
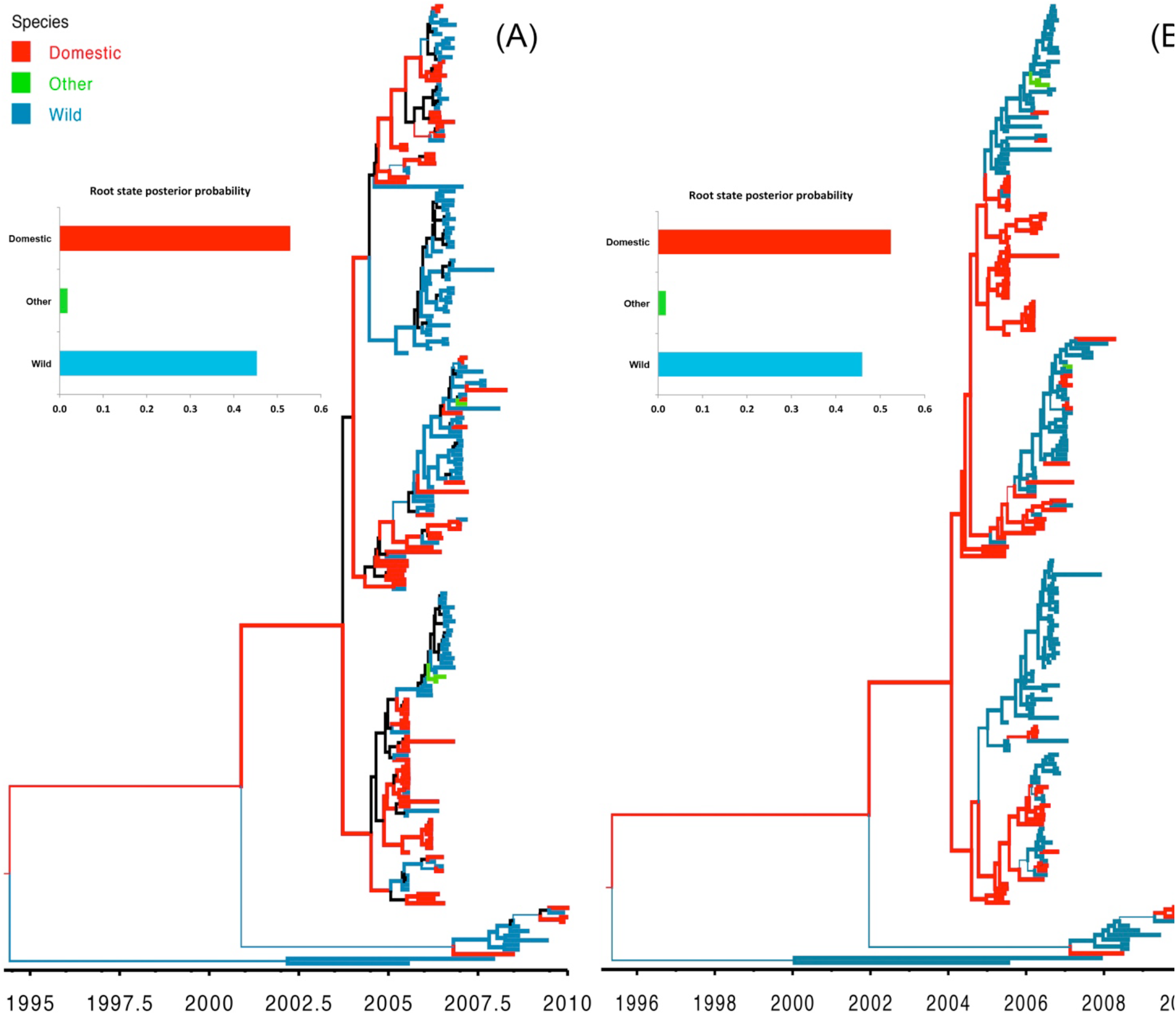
Maximum clade credibility (MCC) trees for H5N1 HPAIV hemagglutinin (HA) and neauraminidase (NA) gene regions (A and B, respectively). Branch lengths are rendered proportional to absolute time (see timescales), the branches are colored according to the most probable host type (wild birds, domestic birds, or other), and the branch thickness is proportional to the posterior probability of the inferred host state. Branches where the host state is uncertain (where the posterior probability of any hosts <0.5) are colored black. The posterior probabilities for the ancestral host states are shown in the upper left panel for each tree.

**Table 1.** Bayes factor (BF) tests for non-zero transition rates between host types for each gene region (HA, NA). BF values > 6 indicate significant rates of exchange between host types.

| Gene | BF | From | To |
| --- | --- | --- | --- |
| HA | 11049.7 | Wild | Other |
| HA | 11049.7 | Other | Wild |
| HA | 171.4 | Domestic | Wild |
| NA | 11049.7 | Wild | Other |
| NA | 11049.7 | Other | Wild |
| NA | 183.0 | Domestic | Wild |

Our divergence-time estimates (Figure 2) suggest that H5N1 HPAIV originated and experienced reassortment among hosts between 1995–1996 (Figure S1). These results are consistent with the frequent reports of highly reassorted H5N1 HPAIVs introductions into eastern and southeastern Asia since the emergence of the ancestral strain in 1996 (A/goose/ Guangdong /1/96) [5, 36].

Results of our discrete phylodynamic analyses indicate that Russia, Romania, and Germany were important epicenters of the H5N1 HPAIV epidemic in Europe. Specifically, our results suggest that the virus first originated and accumulated in Russia, with subsequent significant dispersal rates out of Russia into Romania (Figures 3–5, S2). Later, significant dispersal rates occurred between Romania and Germany. In fact, exchange between Russia-Romania and Romania-Germany represent the most significant dispersal rates of the virus during the course of the epidemic (Figure 5, Table S3). Finally, in late 2005 and early 2006 the virus spread independently from Romania and Germany, respectively into other areas of the EU (Figure S3); a scenario that is consistent with the results of previous studies [1, 14–16, 30].

**Figure 3.**
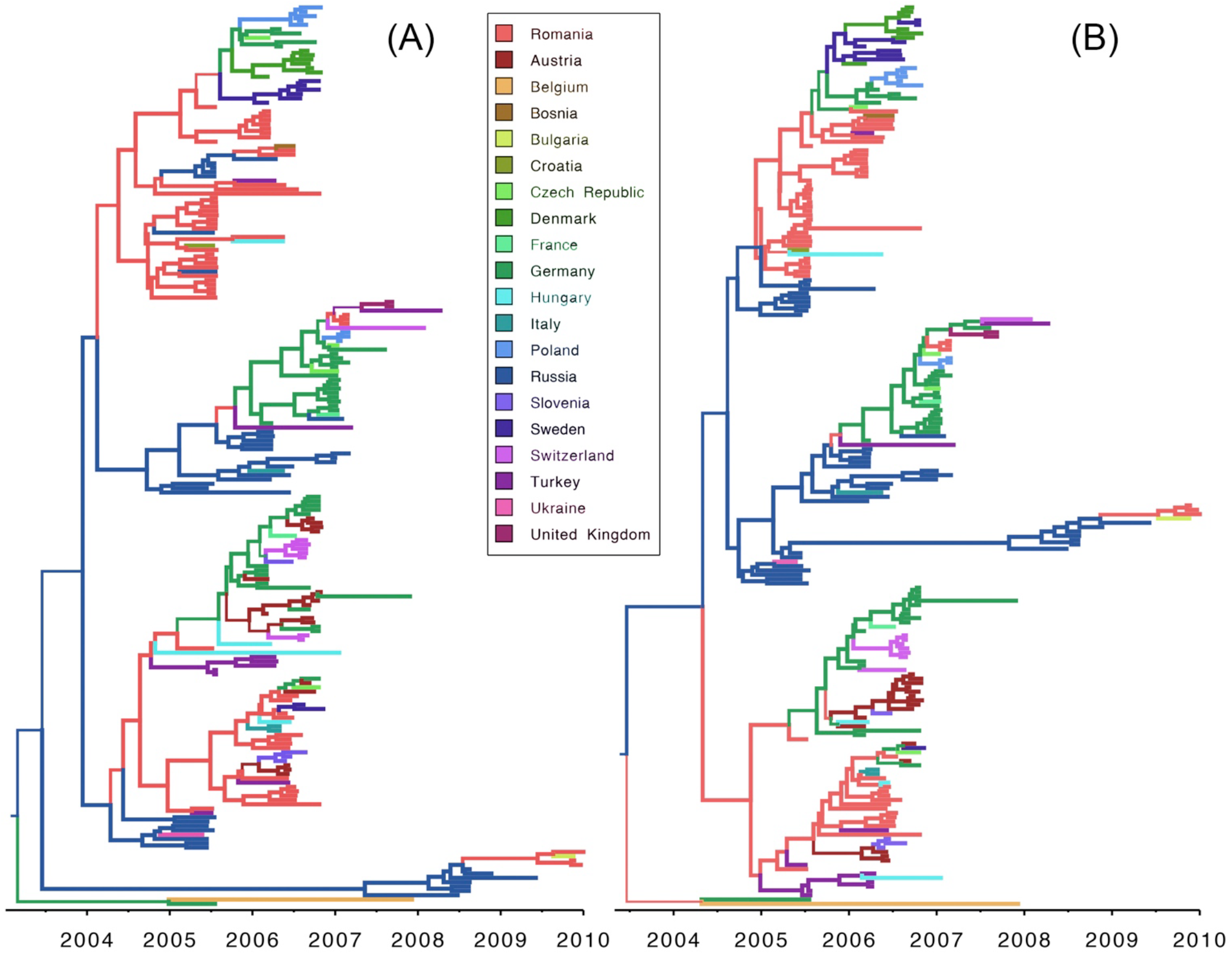
Maximum clade credibility (MCC) tree for H5N1 HPAIV hemagglutinin (HA) and neauraminidase (NA) gene regions. Branch lengths are rendered proportional to absolute time (see timescales), the branches are colored according to the most probable geographic location, and the branch thickness is proportional to the posterior probability of the inferred area (see inset legends). Color gradients along branches reflect the inferred changes in the posterior probability of the corresponding areas.

**Figure 4.**
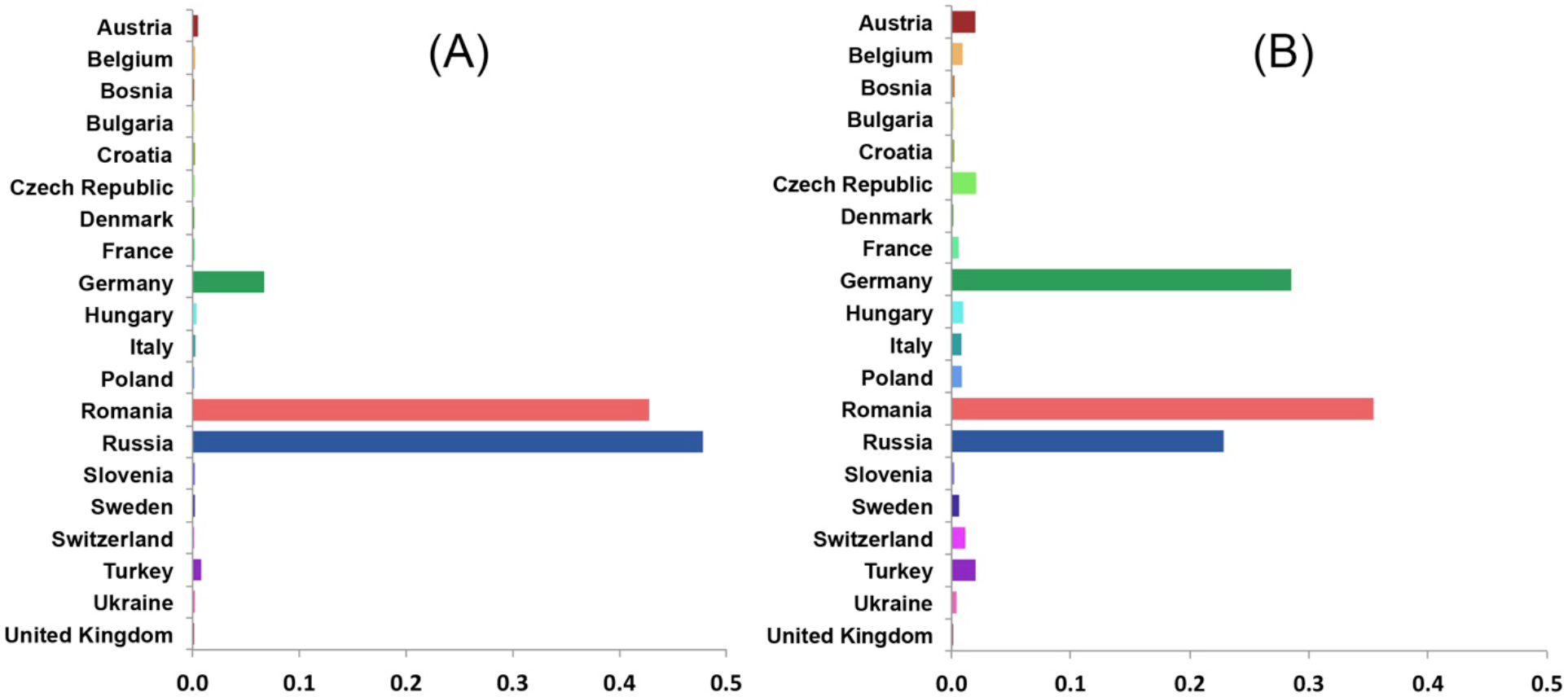
Posterior probabilities of ancestral areas of H5N1 HPAIV hemagglutinin (HA; left) and neauraminidase (NA; right) gene regions collected in Europe and the Russia.

**Figure 5.**
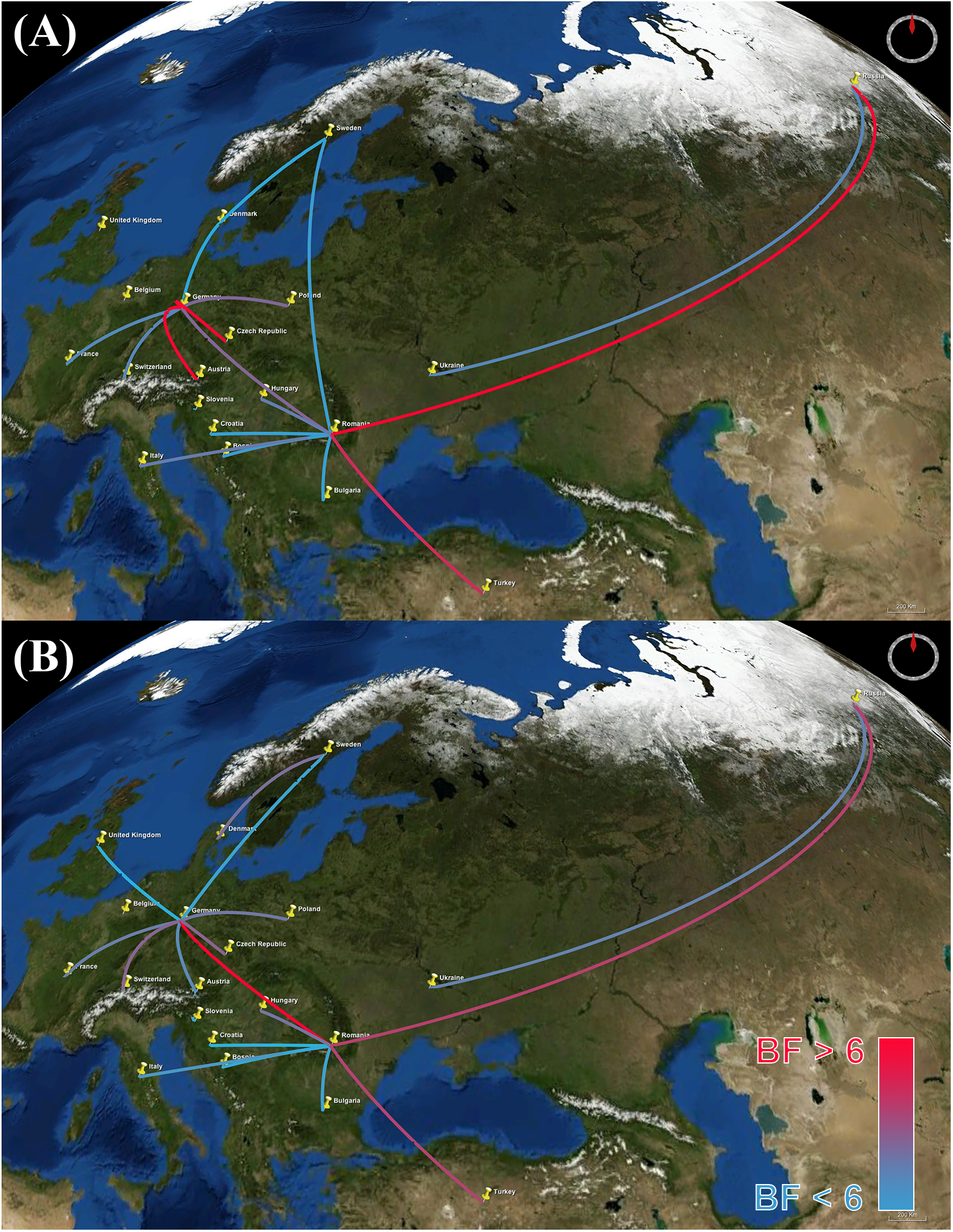
Bayes factor (BF) test for significant non-zero dispersal rates in H5N1 HPAIV. Only rates supported by a BF greater than six are indicated. The color gradient of lines correspond to the probability of the inferred dispersal routes; blue lines and red lines indicate relatively weak and strong support, respectively. The maps are based on satellite pictures available in Google Earth (http://earth.google.com). A: HA gene. B: NA gene. The maps are based on satellite images sourced from the NASA World Wind Java SDK (http://worldwind.arc.nasa.gov/java/).

The inferred geographic expansion of the virus is consistent with our estimates of the viral effective population size; the geographic expanse and the genetic diversity of the virus increased between 2005 and 2006, and subsequently declined between 2007 and 2010 (Figure 1 and S3). The geographic expansion of the virus entailed two major waves. The first wave began in early 2005 from Russia, intensified in Romania, and resulted in the spread of the virus into Germany, Ukraine, and Turkey. The second wave began in early 2006, also commencing in Russia, intensified in both Romania and Germany, and resulted in the spread of the virus into Switzerland, Czech Republic, Denmark, and Sweden [14, 15, 17, 30, 37]. Additionally, a minor wave in 2007 emerged and intensified within Germany, and resulted in the spread of the virus into France, Hungary, and Poland [15].

The results of our phylodynamic analyses generally appear to be robust (Table 2). The significance (p-value <0.001) of the observed association-index (AI) values and their 95% credible intervals for both genes indicate a strong relationship between the inferred phylogeny of the H5N1 virus and the discrete traits (geographic area and host type), and also indicate that both host type and geography played important roles in the transmission of the virus. By contrast, the Kullback-Leibler (KL) values for the discrete host-type models were fairly small for both genes (suggesting relatively poor model fit), whereas the KL values for the geographic models were large, indicating good fit the between model and the geographic data [27].

**Table 2.**
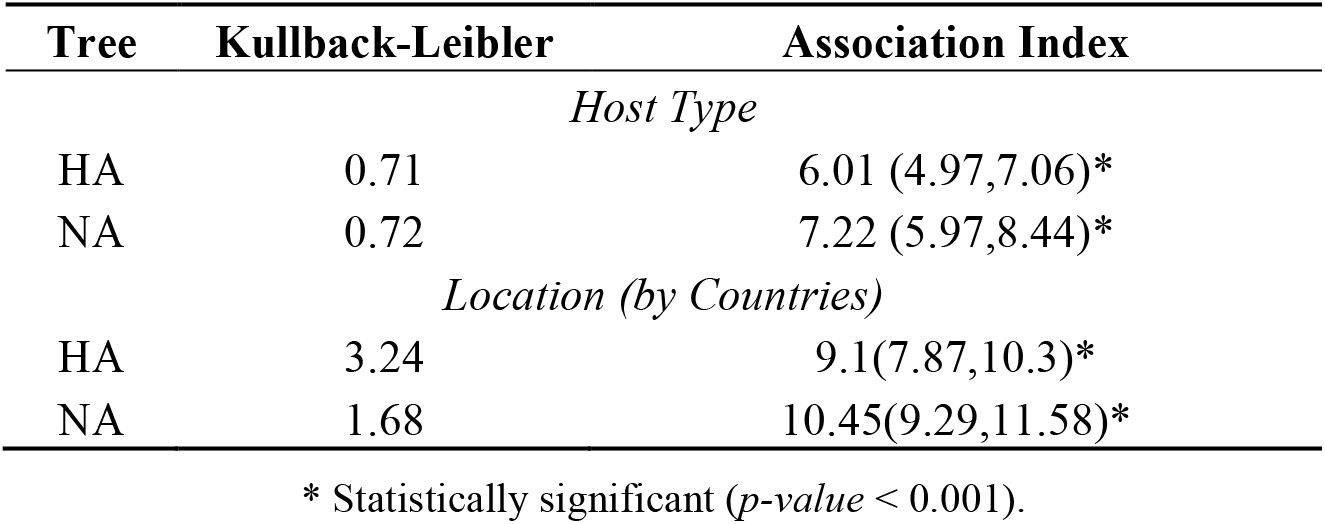
Kullback-Leibler divergence and Association Index statistics assessing the fit of the viral data to the discrete phylodynamic models.

Although generally robust, the results of this study are based on analyses that necessarily entailed several compromises, including: (1) imprecise geographic information on the isolation of H5N1 sequences; and; (2) incomplete and possibly biased sampling of H5N1 isolates. First, precise latitude/longitude coordinates were unavailable for 65.8% of the sequences used in this study; instead only the country of isolation was recorded. In these cases, we used the centroid of the country as a proxy for the location, which is apt to depart substantially from the true location in most cases, which may bias our inferences of the phylogeographic history of the H5N1 virus. This sampling issue precludes our use of the continuous phylodynamic model described by Lemey et al [38], as these models require more precise information on the locations of the viral samples. Unfortunately, this information in typically not recorded for publically available viral sequence data in GenBank or disease databases. The greatest obstacle for applying continuous phylodynamic models to the H5N1 system is therefore mainly an issue of data confidentiality. The impact of incomplete spatial metadata on the performance of phylogeographic models has been discussed elsewhere [39, 40]. Inferences under the phylodynamic models—like all statistical inference methods—assume that we have either a complete or random sample of sequence data. In the present case, this requires that the H5N1 sequences were collected randomly with respect to time (between ∼1990 and 2010), geographic location, and host type. Myriad practical considerations almost certainly result in very strongly biased samples, and the impact of these departures from random sampling on the estimates are difficult to quantify. Although certainly incomplete, and almost certainly non-random, our study is based on all available sequence data for the HA and NA gene regions associated with the H5N1 epidemic in the EU during 2005–2010, and therefore reflects our best understanding based of the available data.

Phylodynamic analysis has not yet been widely embraced as a resource by public health agencies to support the design of disease-surveillance strategies. For example, most of the published literature on avian influenza—including disease summary reports and peer-reviewed scientific studies—upon which global disease agencies base their decision-making process primarily include numerical summaries of reported cases in each country, risk maps based purely on spatial and temporal distribution of the disease, and conventional phylogenetic analysis of sequenced isolates from these outbreaks [13–18, 22– 24, 42]. The 2009 AI global pandemic demonstrates the ability of phylodynamic analyses to provide novel insights for decisions regarding animal and public health [42]. Our phylodynamic analyses of an H5N1 HPAIV sequence dataset and associated metadata provides insights on the origin of the outbreak in Europe (*e.g.*, the ancestral location or host type) and the temporal and the spatial progression of this epidemic. These inferences could be used to inform prevention and control measures to block the virus at the source (*e.g.*, high-risk geographical areas or bird species), which can limit the spread of the virus both to the domestic poultry population or to uninfected geographic areas. Furthermore, the ability to infer and predict routes of viral transmission has clear implications for informing the selection of appropriate strains for vaccine production to more effectively control new reassortant strains of AIV in future epidemics. Accordingly, incorporating phylodynamic analyses as a standard tool for the molecular surveillance of AI might support the development of more effective (and cost effective) policy decisions for the control of this virus in high-risk regions.

## 3. Conclusions

The purpose of this study was to illustrate the potential of a Bayesian phylodynamic approach to inform AI surveillance and control programs and associated decision-making processes. Our results indicate that the peak in viral genetic diversity between 2005–2006 coincided with its geographic spread from Russia into Romania and Germany, and throughout the rest of the EU. This expansion was followed by a drastic decline in viral genetic diversity, which corresponds with the epidemic decline in most EU member states. Our findings corroborate previous conclusions that wild birds were the most important vectors for the spread of the disease. However, our analyses indicate that domestic (rather than wild) birds were the source of the epidemic, and that viral reassortment most likely occurred in domestic species. Finally, our results provide further demonstration of the potential of the Bayesian phylodynamic approach to effectively study the dynamics of disease origin and spread, which promises to maximize the utility of molecular sequence data for the surveillance and control of emerging diseases. These results will contribute to the design of surveillance and control strategies for H5N1 HPAIV in Europe and provide a methodological framework for molecular epidemiological modeling of emerging novel HPAIVs at regional, national, and international scales. This work can also assist global and European disease agencies in their effort to improve the efficiency of their surveillance systems by sharing and improving their genomic databases in the course of emergence of new AIVs.

## 4. Material and Methods

### 4.1 Sequence data

Our dataset comprised complete (or nearly complete) H5N1 HPAIV haemagglutinin (HA) and neuraminidase (NA) gene sequences isolated in Europe and the Russia between May, 2005 and June, 2010, including information on the date, location, and host from which the sequences were isolated. These data were obtained from the Global Initiative on Sharing All influenza data (GISAID) public database (http://platform.gisaid.org), with the exception of sequences from Romania, which have been described elsewhere [46]. Laboratory recombinants or highly cultured sequences were excluded from the dataset. The length of the HA and NA gene regions ranged from 1621–1803 and 1211–1458 bp, respectively. Our dataset comprised a total of 347 sequences, of which 277 (79.8%) were from Europe and Russia, and 70 (20.2%) were from Asian and African countries that were included to root the phylogenetic tree (Table S1). Furthermore, we cross-matched each retrieved sequence with the NCBI Influenza Virus Resource (IVR) (http://www.ncbi.nlm.nih.gov/genomes/FLU/FLU.html) database to obtain information on the date, location, and host of origin of the sample. Because both IVR and GISAID databases do not report the exact longitude and latitude (and occasionally the actual date of collection), we consulted the metadata field for each sequence isolate to approximate the location and date of collection, as well as to specify the host species. We identified metadata for each sequence by comparing strain names between databases (*e.g.*, A/swan/Germany/6/2007), and, when available, we cross-matched information with the corresponding publication [11, 47–52]. We specified the location for each sample (latitude-longitude) using the centroid of the country from which samples were isolated. We converted the collection date for each sequence into fractional years (decimal days) in order to estimate divergence times. If only the year of isolation was available (1.2% of the sequences), we specified the ages as the mid-point of the corresponding year. Similarly, if only the month of isolation was available (10.4% of the sequences), we recorded the date as the mid-point of the corresponding month. Finally, we classified host species as either *wild birds*, *domestic birds*, or *other* (non-avian hosts). Table S2 summarizes the profile of 277 H5N1 HPAIV sequences isolated from reported outbreaks in Europe and the Russia between May 2005, and June 2010. Finally, an epidemic curve was plotted using approximately 1130 outbreak reports retrieved from the Food and Agriculture Organization of the United Nation (FAO) Global Animal Disease Information System EMPRES-i (http://empres-i.fao.org/). The outbreak data included dates and locations of detected cases in domesticated poultry and wild birds in Europe and Russia between 2005 and 2010.

### 4.2 Preliminary phylogenetic analysis

We aligned the H5N1 HPAIV HA and NA gene regions using MUSCLE version 3.8 [43]. The resulting alignments were adjusted manually to ensure that these protein-coding gene regions remained in frame (assessed by amino-acid translation using Mesquite version 3.01) [44]. We removed all sequences (25%) with 100% nucleotide identity and both gene regions were found to be free of homologous recombination using Recombination Detection Program version 3 (RDP3) [45]. For each gene region, we selected a mixed substitution model (assignment of data partitions to substitution models) by first defining six data subsets (the codon positions of each gene region), and then selecting among mixed-substitution model using the Bayesian Information Criterion (BIC) implemented in Partition-Finder v 1.1 [46]. Finally, we assessed the degree of topological (in)congruence between the two gene regions by performing maximum-likelihood estimates of the phylogeny for the individual gene regions under the selected mixed-substitution model; these analyses entailed 100 non-parametric bootstrap replicate searches of each gene region using RAxML version 8 [47].

### 4.3 Divergence-time estimation

We estimated divergence times for the H5N1 HPAIV sequence dataset using the (relaxed) molecu-lar-clock models implemented in BEAST v 1.8 [48]. We used the mixed-substitution model for each gene region identified in the preliminary analyses detailed above. We assessed the fit of the sequence data to three branch-rate prior models: (1) a strict molecular clock model (which assumes that substitution rates are stochastically constant across branches of the tree); (2) the uncorrelated exponential relaxed clock (UCED) model, which assumes that substitution rates on adjacent branches are sampled from a shared exponential distribution, and; (3) the uncorrelated lognormal relaxed clock (UCLN) model, which assumes that substitution rates on adjacent branches are drawn from a shared lognormal distribution. We assessed the fit of the sequence data to these three branch-rate models by comparing their corresponding marginal likelihoods, which were estimated using Tracer version 1.6 [48, 49]. Bayes factor (BF) comparisons indicated that the UCED branch-rate model provided the best fit for both HA and NA gene regions (Bayes factor >25 for the log marginal likelihood). Estimation of divergence times also requires a node-age model; we assumed a Bayesian skyline coalescent tree prior, which allowed us to estimate changes in the effective population size through time [50]. Isolation dates of the sequences were used to calibrate divergence-time estimates.

We approximated the joint posterior probability density of these model parameters using the Markov Chain Monte Carlo (MCMC) algorithms implemented in BEAST v 1.8. Each MCMC simulation was run for 1x10^8^ cycles, and thinned by sampling every 10,000 cycle. We performed four replicate MCMC simulations to aid in assessing performance of the simulations. We assessed MCMC convergence by comparing the parameter estimates from independent analyses for each parameter, ensuring that the estimates of the marginal posterior probability densities were effectively identical for the four independent chains. We also assessed convergence by the estimating effective sample sizes (ESS) for each parameter using Tracer, and assessed mixing based on the acceptance ratios. Those evaluations suggested that the MCMC algorithms provided a reliable approximation of the posterior probability density, and suggested that the first 10% of the samples (the “burn-in”) should be discarded. We summarized posterior results as a maximum clade credibility (MCC) tree using TreeAnnotator. We then generated a Bayesian skyride plot to infer the population demographics of H5N1 HPAIV HA and NA gene segments in Europe and the Russia. We plotted the inferred effective population size of the virus between 2005–2010 in terms of relative genetic diversity (*N*_*e*_*T*), where *N*_*e*_ is the effective population size and *T* the generation time [51].

### 4.4 Estimation of geographic history under the discrete phylodynamic model

We incorporated information on the geographic locations where sequences were isolated to describe the spatial dynamics of viral epidemics following a procedure described by Lemey *et al.* [27]. Briefly, the approach jointly estimates phylogeny and history of discrete traits in a Bayesian statistical framework, which enables inference of the timing of viral dispersal patterns while accommodating phylogenetic uncertainty. Here, the discrete states are geographic areas, and the goal is to estimate the history of viral migration (transitions) between these areas through time. This is achieved using a model averaging approach (based on Bayesian stochastic search variable selection; BSSVS) to describe the spatial evolution of H5N1 HPAIV epidemic. We modeled the geographic transition of the H5N1 HPAIV between areas as discrete states under a continuous-time Markov model comprising a number of nonzero transition rates that were identified by means of BSSVS. We assessed the fit of the H5N1 HPAIV data to a number of candidate discrete phylogeographic models (Table S4), including both symmetric and asymmetric discrete-trait models with irreversible and reversible transitions, respectively (using a mean-one exponential prior for the rate parameters). Accordingly, our analyses were based on a composite phylogenetic model that comprised: (1) the previously selected mixed-substitution model; (2) the strict, UCLN, UCED branch-rate models to describe variation in substitution rate across branches; (3) the coalescent Gaussian Markov Random field (GMRF) Bayesian Skyride model as a prior on the node times in the tree, and; (4) the symmetric and asymmetric discrete-state phylodynamic models to describe the history of viral migration between discrete geographic areas. Accordingly, we explored a total of six candidate composite phylodynamic models (the symmetric and asymmetric variants of the geographic model and the three branch-rate models).

We assessed the relative fit of the six candidate phylodynamic models to the HA and NA sequence datasets using Bayes factors. To this end, we first estimated the marginal likelihood for each of the six candidate phylodynamic models from the resulting posterior samples using the posterior simulation-based analogue of Akaike’s information criterion (AICm) [52]. Bael et al. [53] have recently shown the AICm to provide more reliable estimates of the marginal likelihood than the harmonic-mean estimator. We then used the marginal-likelihood estimates to compute Bayes factors to select among the candidate models. The UCED branch-rate model was again preferred for both HA and NA gene segments (BF > 25). Bayes factor (BF) comparisons indicated that the symmetric UCED branch-rate model with irreversible transitions provided the best fit for both HA and NA gene regions (Bayes factor >25 for the log marginal likelihood). The posterior probability distribution of trees sampled under the preferred model was summarized as an MCC consensus tree with posterior probabilities of geographic areas plotted at interior nodes using FigTree version 1.4 [54]. We used SPREAD version 1.0.6 [55] to assess the strength of transition rates between discrete geographic areas, using a BF cutoff = 6 to identify non-zero transition rates. Finally, we generated a keyhole markup language (KML) file to visualize the geographic migration of the virus.

### 4.5 Exploring the evolution of H5N1 HPAIV host infection

We modeled the evolutionary movement of H5N1 HPAIV within and between host types (domestic birds, wild birds, other). To this end, we again adopted a discrete-trait model, where the discrete states in this case correspond to the host type, where the objective is to infer the history of H5N1 HPAIV migration between hosts through time. The number of non-zero transition rates in the model was again estimated using BSSVS. The relative strength of transition rates (*e.g.*, wild → domestic) was estimated using Bayes factors. Following [64], we estimated the ancestral states (host type) at internal nodes of the tree under a composite phylogenetic model that included: (1) the previously selected mixed-substitution model to describe evolution of nucleotides over the tree with branch lengths; (2) the preferred uncorrelated lognormal model to describe the variation of substitution rates across branches of the tree [56]; (3) a constant population-size coalescent model to describe the temporal distribution of node heights in the phylogeny, and; (4) the asymmetric discrete-state phylodynamic model to describe the history of viral migration between host types as a continuous-time Markov process (with a number of non-zero transition rates determined by the BSSVS procedure, using a uniform prior with mean = 1 for the rate parameters). We summarized the posterior probability distribution of parameter estimates as an MCC consensus tree with the posterior probability of host state mapped to interior nodes of the tree using FigTree. Additionally, we assessed the strength of transition rates between states (hosts) by means of Bayes factors using SPREAD with a cutoff of six to identify non-zero transition rates. Use of the asymmetric discrete trait model allowed us to assess the strength of directionality between states (*e.g.*, wild → domestic and domestic → wild bird).

### 4.6 Assessing uncertainty in discrete-trait mappings and association statistics

We used the Kullback-Leibler (KL) divergence statistic [57] to accommodate phylogenetic uncertainty in the discrete-trait estimates (for host type and geographic location). The KL divergence measures the departure between prior probability distribution and the corresponding posterior probability distribution for a given parameter. The premise is simple: the posterior probability distribution inferred for a given parameter is the updated version of prior probability distribution—it is updated by the information in the data via the likelihood function. Accordingly, if the data contain little information regarding the value of a parameter, its posterior probability distribution will closely resemble the corresponding prior probability distribution, and the KL divergence between these two probability distributions will therefore be small [27]. We calculated the KL divergence for each selected posterior distribution of trees using a function written by Razavi [58] implemented in Matlab v 2013a [59]. The Association Index (AI) was calculated using Bayesian Tip-Significance Testing (BaTS) software version 1.0 to test the hypothesis that a taxon with a given trait (host type or geographic location) are more likely to share traits with adjoining taxa than that expected by chance [60].

## Acknowledgments

We are grateful to Matthew L. Scotch for assistance with Matlab. This research was supported by National Science Foundation (NSF) grants DEB-0842181, DEB-0919529, DBI-1356737, and DEB-1457835 awarded to B.R.M.

## Author Contributions

M.A. and B.R.M. conceived and designed the study. M.A. and A.P. analyzed the data. B.R.M. provided advise regarding the analyses. M.A., B.R.M. and A.P. wrote the paper.

## Conflicts of Interest

The authors declare no conflict of interest.

## Supplementary Materials

**Table S1.** Bayes factor test for non-zero rates for location (country) state model for both HA and NA genes. Large BF values indicate most likely routes of dispersal between country’s location states.

| Gene | BF | Between |  |
| --- | --- | --- | --- |
| HA | 77823.6 | Russia | Romania |
| HA | 77823.6 | Germany | Austria |
| HA | 77823.6 | Germany | Czech Republic |
| HA | 8639.4 | Turkey | Romania |
| HA | 317.0 | Romania | Germany |
| HA | 274.4 | Poland | Germany |
| HA | 90.6 | Romania | Hungary |
| HA | 87.4 | Romania | Italy |
| HA | 48.5 | Ukraine | Russia |
| HA | 46.4 | Switzerland | Germany |
| HA | 43.4 | Germany | France |
| HA | 16.0 | Romania | Bosnia |
| HA | 13.9 | Sweden | Romania |
| HA | 11.9 | Romania | Bulgaria |
| HA | 11.1 | Romania | Croatia |
| HA | 9.8 | Slovenia | Austria |
| HA | 7.7 | Sweden | Denmark |
| HA | 7.0 | Germany | Denmark |
| NA | 77823.6 | Romania | Germany |
| NA | 2984.9 | Russia | Romania |
| NA | 2502.1 | Turkey | Romania |
| NA | 441.2 | Germany | Czech Republic |
| NA | 409.8 | Switzerland | Germany |
| NA | 240.0 | Sweden | Denmark |
| NA | 190.9 | Romania | Hungary |
| NA | 149.2 | Poland | Germany |
| NA | 108.2 | Germany | France |
| NA | 87.0 | Ukraine | Russia |
| NA | 57.9 | Germany | Austria |
| NA | 27.5 | Romania | Italy |
| NA | 16.3 | Romania | Bulgaria |
| NA | 15.4 | Slovenia | Austria |
| NA | 15.3 | Sweden | Germany |
| NA | 11.1 | Romania | Bosnia |
| NA | 7.6 | United Kingdom | Germany |
| NA | 6.9 | Romania | Croatia |

Bayes factors (BF) > 6 indicate significant non-zero dispersal

**Table S2.**
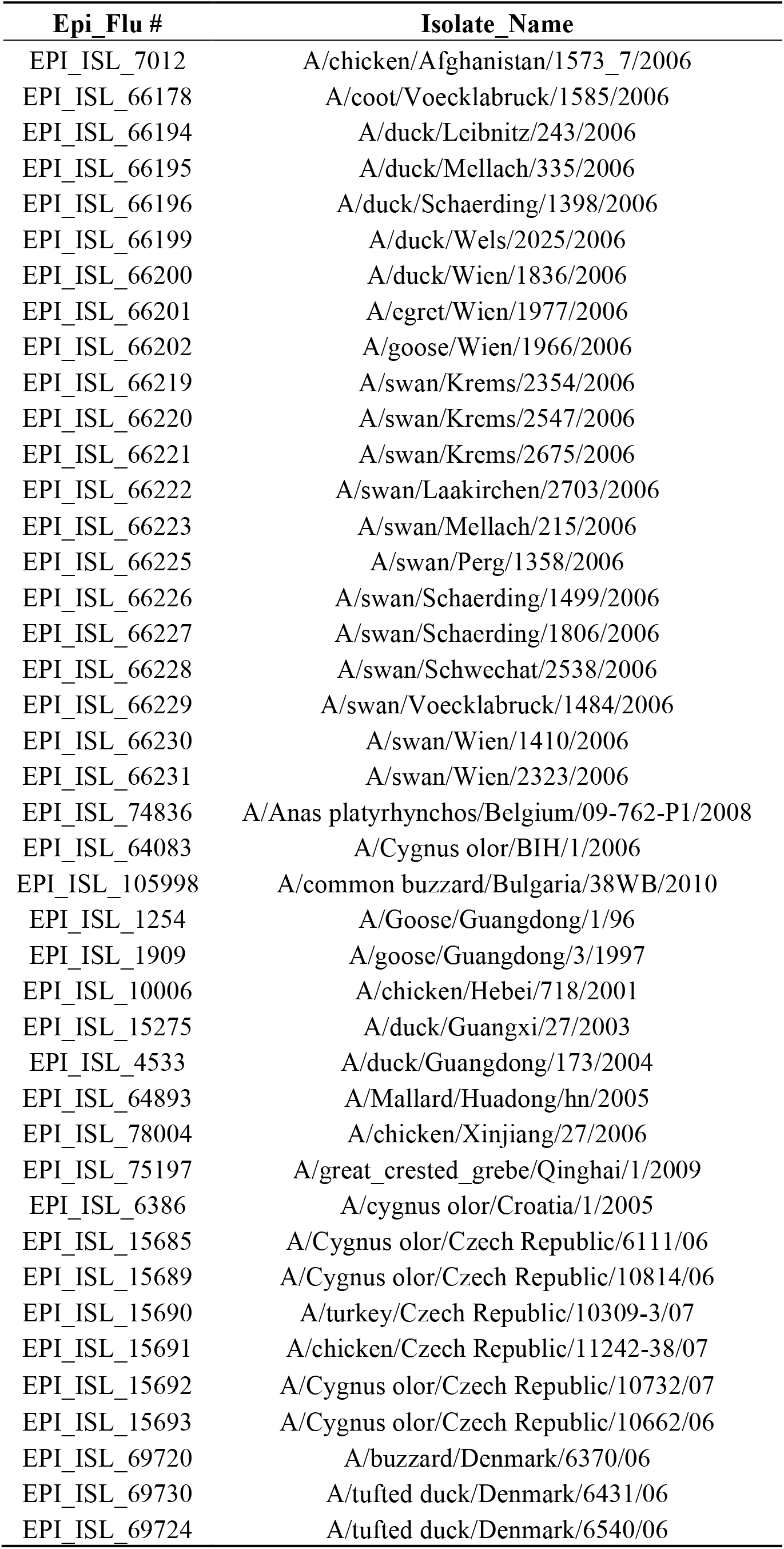
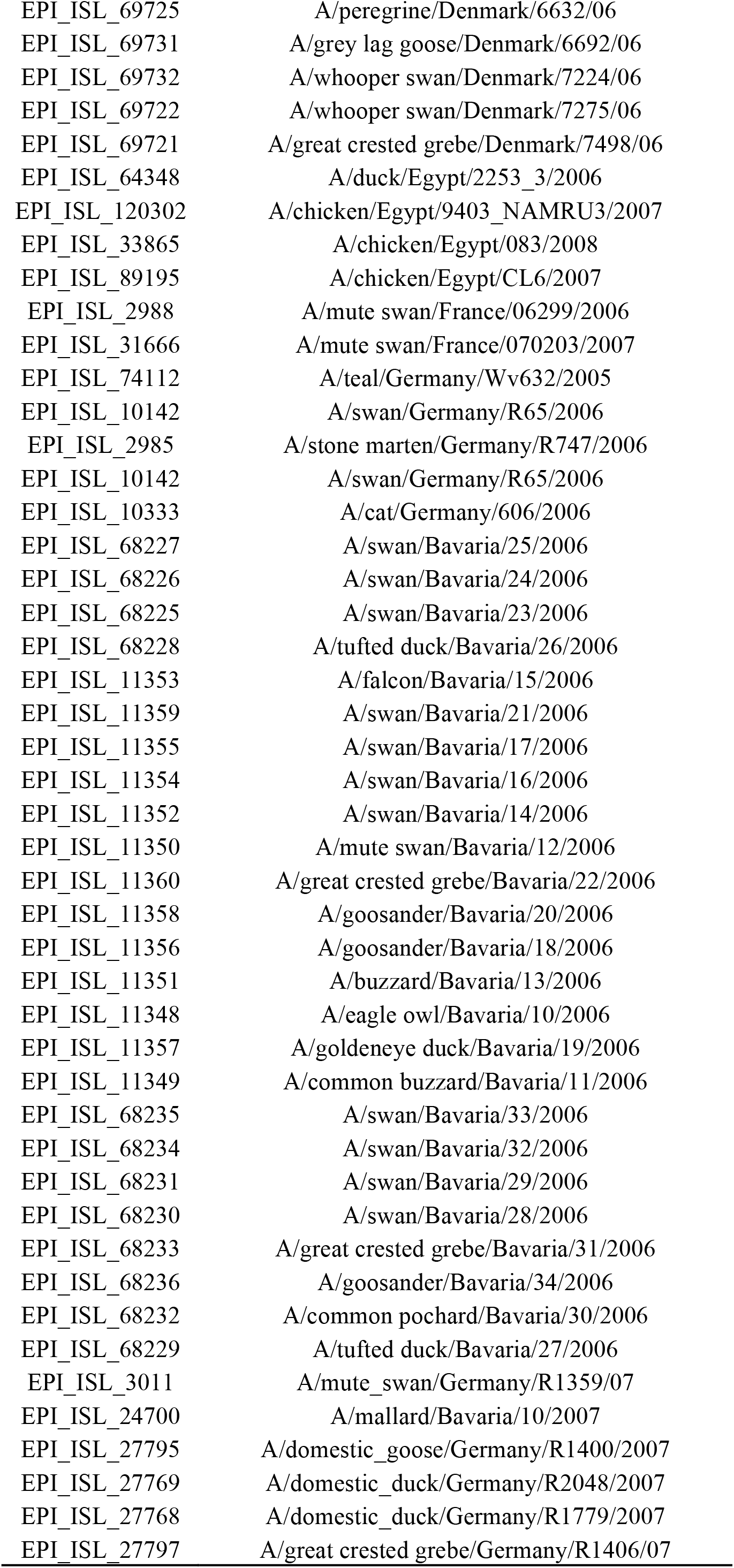
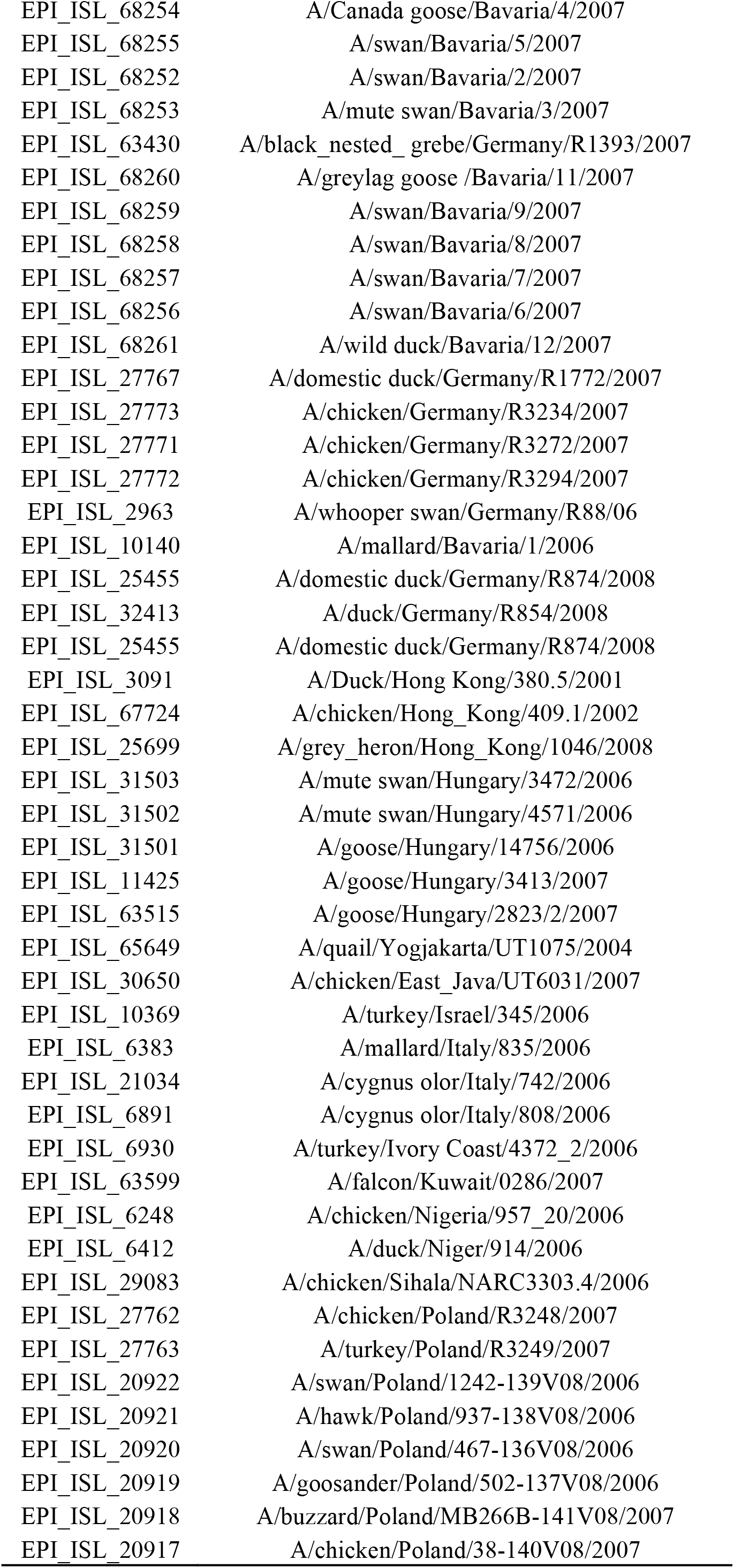
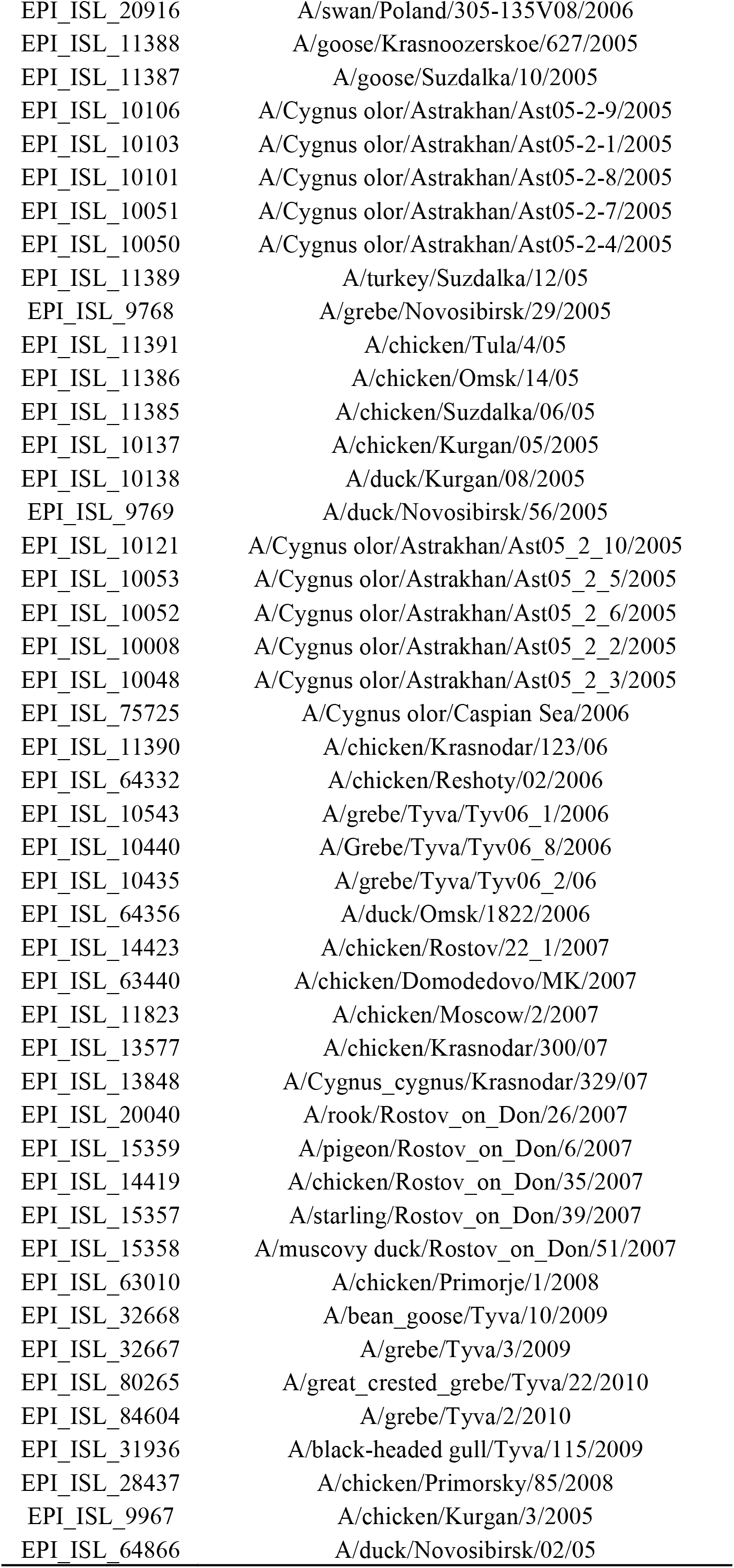
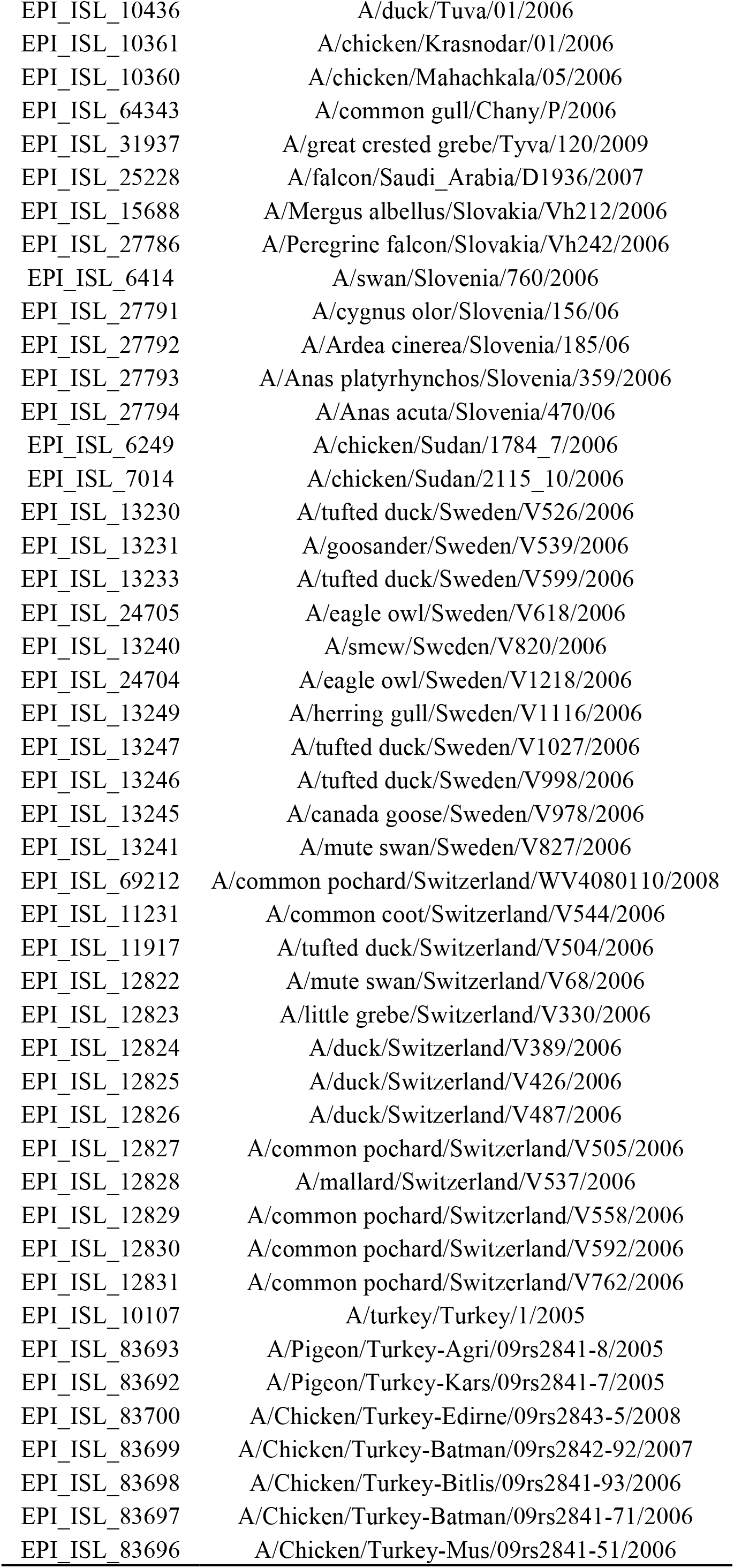
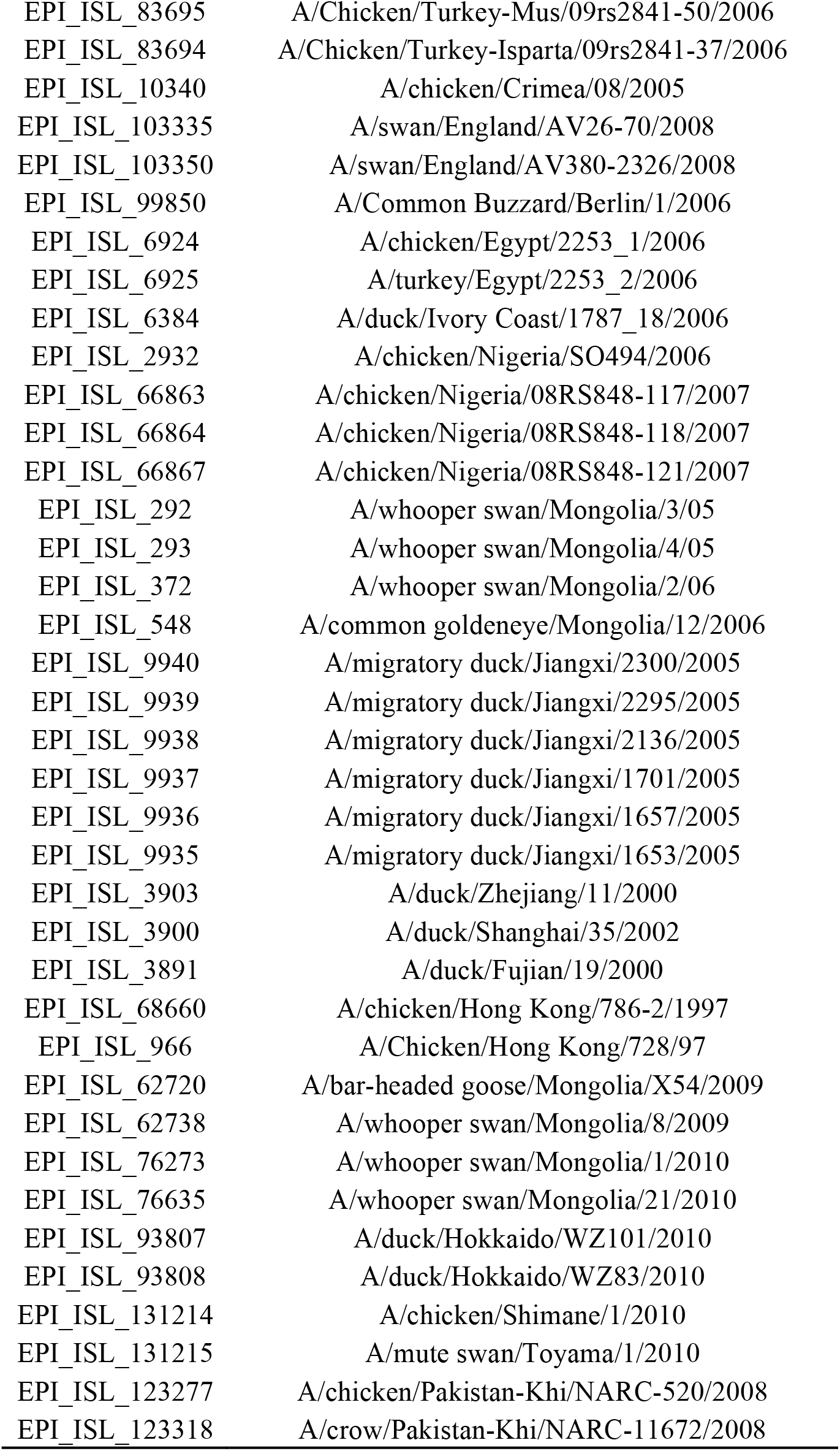
GISAID database Accession numbers and isolate names matched with other public databases used in this study for both HA and NA gene segments, with the exception of sequences isolated from Romania.

| Epi_Flu # | Isolate_Name |
| --- | --- |
| EPI_ISL_7012 | A/chicken/Afghanistan/1573_7/2006 |
| EPI_ISL_66178 | A/coot/Voecklabruck/1585/2006 |
| EPI_ISL_66194 | A/duck/Leibnitz/243/2006 |
| EPI_ISL_66195 | A/duck/Mellach/335/2006 |
| EPI_ISL_66196 | A/duck/Schaerding/1398/2006 |
| EPI_ISL_66199 | A/duck/Wels/2025/2006 |
| EPI_ISL_66200 | A/duck/Wien/1836/2006 |
| EPI_ISL_66201 | A/egret/Wien/1977/2006 |
| EPI_ISL_66202 | A/goose/Wien/1966/2006 |
| EPI_ISL_66219 | A/swan/Krems/2354/2006 |
| EPI_ISL_66220 | A/swan/Krems/2547/2006 |
| EPI_ISL_66221 | A/swan/Krems/2675/2006 |
| EPI_ISL_66222 | A/swan/Laakirchen/2703/2006 |
| EPI_ISL_66223 | A/swan/Mellach/215/2006 |
| EPI_ISL_66225 | A/swan/Perg/1358/2006 |
| EPI_ISL_66226 | A/swan/Schaerding/1499/2006 |
| EPI_ISL_66227 | A/swan/Schaerding/1806/2006 |
| EPI_ISL_66228 | A/swan/Schwechat/2538/2006 |
| EPI_ISL_66229 | A/swan/Voecklabruck/1484/2006 |
| EPI_ISL_66230 | A/swan/Wien/1410/2006 |
| EPI_ISL_66231 | A/swan/Wien/2323/2006 |
| EPI_ISL_74836 | A/Anas platyrhynchos/Belgium/09-762-P1/2008 |
| EPI_ISL_64083 | A/Cygnus olor/BIH/1/2006 |
| EPI_ISL_105998 | A/common buzzard/Bulgaria/38WB/2010 |
| EPI_ISL_1254 | A/Goose/Guangdong/1/96 |
| EPI_ISL_1909 | A/goose/Guangdong/3/1997 |
| EPI_ISL_10006 | A/chicken/Hebei/718/2001 |
| EPI_ISL_15275 | A/duck/Guangxi/27/2003 |
| EPI_ISL_4533 | A/duck/Guangdong/173/2004 |
| EPI_ISL_64893 | A/Mallard/Huadong/hn/2005 |
| EPI_ISL_78004 | A/chicken/Xinjiang/27/2006 |
| EPI_ISL_75197 | A/great_crested_grebe/Qinghai/1/2009 |
| EPI_ISL_6386 | A/cygnus olor/Croatia/1/2005 |
| EPI_ISL_15685 | A/Cygnus olor/Czech Republic/6111/06 |
| EPI_ISL_15689 | A/Cygnus olor/Czech Republic/10814/06 |
| EPI_ISL_15690 | A/turkey/Czech Republic/10309-3/07 |
| EPI_ISL_15691 | A/chicken/Czech Republic/11242-38/07 |
| EPI_ISL_15692 | A/Cygnus olor/Czech Republic/10732/07 |
| EPI_ISL_15693 | A/Cygnus olor/Czech Republic/10662/06 |
| EPI_ISL_69720 | A/buzzard/Denmark/6370/06 |
| EPI_ISL_69730 | A/tufted duck/Denmark/6431/06 |
| EPI_ISL_69724 | A/tufted duck/Denmark/6540/06 |
| EPI_ISL_69725 | A/peregrine/Denmark/6632/06 |
| EPI_ISL_69731 | A/grey lag goose/Denmark/6692/06 |
| EPI_ISL_69732 | A/whooper swan/Denmark/7224/06 |
| EPI_ISL_69722 | A/whooper swan/Denmark/7275/06 |
| EPI_ISL_69721 | A/great crested grebe/Denmark/7498/06 |
| EPI_ISL_64348 | A/duck/Egypt/2253_3/2006 |
| EPI_ISL_120302 | A/chicken/Egypt/9403_NAMRU3/2007 |
| EPI_ISL_33865 | A/chicken/Egypt/083/2008 |
| EPI_ISL_89195 | A/chicken/Egypt/CL6/2007 |
| EPI_ISL_2988 | A/mute swan/France/06299/2006 |
| EPI_ISL_31666 | A/mute swan/France/070203/2007 |
| EPI_ISL_74112 | A/teal/Germany/Wv632/2005 |
| EPI_ISL_10142 | A/swan/Germany/R65/2006 |
| EPI_ISL_2985 | A/stone marten/Germany/R747/2006 |
| EPI_ISL_10142 | A/swan/Germany/R65/2006 |
| EPI_ISL_10333 | A/cat/Germany/606/2006 |
| EPI_ISL_68227 | A/swan/Bavaria/25/2006 |
| EPI_ISL_68226 | A/swan/Bavaria/24/2006 |
| EPI_ISL_68225 | A/swan/Bavaria/23/2006 |
| EPI_ISL_68228 | A/tufted duck/Bavaria/26/2006 |
| EPI_ISL_11353 | A/falcon/Bavaria/15/2006 |
| EPI_ISL_11359 | A/swan/Bavaria/21/2006 |
| EPI_ISL_11355 | A/swan/Bavaria/17/2006 |
| EPI_ISL_11354 | A/swan/Bavaria/16/2006 |
| EPI_ISL_11352 | A/swan/Bavaria/14/2006 |
| EPI_ISL_11350 | A/mute swan/Bavaria/12/2006 |
| EPI_ISL_11360 | A/great crested grebe/Bavaria/22/2006 |
| EPI_ISL_11358 | A/goosander/Bavaria/20/2006 |
| EPI_ISL_11356 | A/goosander/Bavaria/18/2006 |
| EPI_ISL_11351 | A/buzzard/Bavaria/13/2006 |
| EPI_ISL_11348 | A/eagle owl/Bavaria/10/2006 |
| EPI_ISL_11357 | A/goldeneye duck/Bavaria/19/2006 |
| EPI_ISL_11349 | A/common buzzard/Bavaria/11/2006 |
| EPI_ISL_68235 | A/swan/Bavaria/33/2006 |
| EPI_ISL_68234 | A/swan/Bavaria/32/2006 |
| EPI_ISL_68231 | A/swan/Bavaria/29/2006 |
| EPI_ISL_68230 | A/swan/Bavaria/28/2006 |
| EPI_ISL_68233 | A/great crested grebe/Bavaria/31/2006 |
| EPI_ISL_68236 | A/goosander/Bavaria/34/2006 |
| EPI_ISL_68232 | A/common pochard/Bavaria/30/2006 |
| EPI_ISL_68229 | A/tufted duck/Bavaria/27/2006 |
| EPI_ISL_3011 | A/mute swan/Germany/R1359/07 |
| EPI_ISL_24700 | A/mallard/Bavaria/10/2007 |
| EPI_ISL_27795 | A/domestic_goose/Germany/R1400/2007 |
| EPI_ISL_27769 | A/domestic_duck/Germany/R2048/2007 |
| EPI_ISL_27768 | A/domestic_duck/Germany/R1779/2007 |
| EPI_ISL_27797 | A/great crested grebe/Germany/R1406/07 |
| EPI_ISL_68254 | A/Canada goose/Bavaria/4/2007 |
| EPI_ISL_68255 | A/swan/Bavaria/5/2007 |
| EPI_ISL_68252 | A/swan/Bavaria/2/2007 |
| EPI_ISL_68253 | A/mute swan/Bavaria/3/2007 |
| EPI_ISL_63430 | A/black_nested_grebe/Germany/R1393/2007 |
| EPI_ISL_68260 | A/greylag goose /Bavaria/11/2007 |
| EPI_ISL_68259 | A/swan/Bavaria/9/2007 |
| EPI_ISL_68258 | A/swan/Bavaria/8/2007 |
| EPI_ISL_68257 | A/swan/Bavaria/7/2007 |
| EPI_ISL_68256 | A/swan/Bavaria/6/2007 |
| EPI_ISL_68261 | A/wild duck/Bavaria/12/2007 |
| EPI_ISL_27767 | A/domestic duck/Germany/R1772/2007 |
| EPI_ISL_27773 | A/chicken/Germany/R3234/2007 |
| EPI_ISL_27771 | A/chicken/Germany/R3272/2007 |
| EPI_ISL_27772 | A/chicken/Germany/R3294/2007 |
| EPI_ISL_2963 | A/whooper swan/Germany/R88/06 |
| EPI_ISL_10140 | A/mallard/Bavaria/1/2006 |
| EPI_ISL_25455 | A/domestic duck/Germany/R874/2008 |
| EPI_ISL_32413 | A/duck/Germany/R854/2008 |
| EPI_ISL_25455 | A/domestic duck/Germany/R874/2008 |
| EPI_ISL_3091 | A/Duck/Hong Kong/380.5/2001 |
| EPI_ISL_67724 | A/chicken/Hong_Kong/409.1/2002 |
| EPI_ISL_25699 | A/grey_heron/Hong_Kong/1046/2008 |
| EPI_ISL_31503 | A/mute swan/Hungary/3472/2006 |
| EPI_ISL_31502 | A/mute swan/Hungary/4571/2006 |
| EPI_ISL_31501 | A/goose/Hungary/14756/2006 |
| EPI_ISL_11425 | A/goose/Hungary/3413/2007 |
| EPI_ISL_63515 | A/goose/Hungary/2823/2/2007 |
| EPI_ISL_65649 | A/quail/Yogyakarta/UT1075/2004 |
| EPI_ISL_30650 | A/chicken/East_Java/UT6031/2007 |
| EPI_ISL_10369 | A/turkey/Israel/345/2006 |
| EPI_ISL_6383 | A/mallard/Italy/835/2006 |
| EPI_ISL_21034 | A/cygnus olor/Italy/742/2006 |
| EPI_ISL_6891 | A/cygnus olor/Italy/808/2006 |
| EPI_ISL_6930 | A/turkey/Ivory Coast/4372_2/2006 |
| EPI_ISL_63599 | A/falcon/Kuwait/0286/2007 |
| EPI_ISL_6248 | A/chicken/Nigeria/957_20/2006 |
| EPI_ISL_6412 | A/duck/Niger/914/2006 |
| EPI_ISL_29083 | A/chicken/Sihala/NARC3303.4/2006 |
| EPI_ISL_27762 | A/chicken/Poland/R3248/2007 |
| EPI_ISL_27763 | A/turkey/Poland/R3249/2007 |
| EPI_ISL_20922 | A/swan/Poland/1242-139V08/2006 |
| EPI_ISL_20921 | A/hawk/Poland/937-138V08/2006 |
| EPI_ISL_20920 | A/swan/Poland/467-136V08/2006 |
| EPI_ISL_20919 | A/goosander/Poland/502-137V08/2006 |
| EPI_ISL_20918 | A/buzzard/Poland/MB266B-141V08/2007 |
| EPI_ISL_20917 | A/chicken/Poland/38-140V08/2007 |
| EPI_ISL_20916 | A/swan/Poland/305-135V08/2006 |
| EPI_ISL_11388 | A/goose/Krasnoozerskoe/627/2005 |
| EPI_ISL_11387 | A/goose/Suzdalka/10/2005 |
| EPI_ISL_10106 | A/Cygnus olor/Astrakhan/Ast05-2-9/2005 |
| EPI_ISL_10103 | A/Cygnus olor/Astrakhan/Ast05-2-1/2005 |
| EPI_ISL_10101 | A/Cygnus olor/Astrakhan/Ast05-2-8/2005 |
| EPI_ISL_10051 | A/Cygnus olor/Astrakhan/Ast05-2-7/2005 |
| EPI_ISL_10050 | A/Cygnus olor/Astrakhan/Ast05-2-4/2005 |
| EPI_ISL_11389 | A/turkey/Suzdalka/12/05 |
| EPI_ISL_9768 | A/grebe/Novosibirsk/29/2005 |
| EPI_ISL_11391 | A/chicken/Tula/4/05 |
| EPI_ISL_11386 | A/chicken/Omsk/14/05 |
| EPI_ISL_11385 | A/chicken/Suzdalka/06/05 |
| EPI_ISL_10137 | A/chicken/Kurgan/05/2005 |
| EPI_ISL_10138 | A/duck/Kurgan/08/2005 |
| EPI_ISL_9769 | A/duck/Novosibirsk/56/2005 |
| EPI_ISL_10121 | A/Cygnus olor/Astrakhan/Ast05_2_10/2005 |
| EPI_ISL_10053 | A/Cygnus olor/Astrakhan/Ast05_2_5/2005 |
| EPI_ISL_10052 | A/Cygnus olor/Astrakhan/Ast05_2_6/2005 |
| EPI_ISL_10008 | A/Cygnus olor/Astrakhan/Ast05_2_2/2005 |
| EPI_ISL_10048 | A/Cygnus olor/Astrakhan/Ast05_2_3/2005 |
| EPI_ISL_75725 | A/Cygnus olor/Caspian Sea/2006 |
| EPI_ISL_11390 | A/chicken/Krasnodar/123/06 |
| EPI_ISL_64332 | A/chicken/Reshoty/02/2006 |
| EPI_ISL_10543 | A/grebe/Tyva/Tyv06_1/2006 |
| EPI_ISL_10440 | A/Grebe/Tyva/Tyv06_8/2006 |
| EPI_ISL_10435 | A/grebe/Tyva/Tyv06_2/06 |
| EPI_ISL_64356 | A/duck/Omsk/1822/2006 |
| EPI_ISL_14423 | A/chicken/Rostov/22_1/2007 |
| EPI_ISL_63440 | A/chicken/Domodedovo/MK/2007 |
| EPI_ISL_11823 | A/chicken/Moscow/2/2007 |
| EPI_ISL_13577 | A/chicken/Krasnodar/300/07 |
| EPI_ISL_13848 | A/Cygnus_cygnus/Krasnodar/329/07 |
| EPI_ISL_20040 | A/rook/Rostov_on_Don/26/2007 |
| EPI_ISL_15359 | A/pigeon/Rostov_on_Don/6/2007 |
| EPI_ISL_14419 | A/chicken/Rostov_on_Don/35/2007 |
| EPI_ISL_15357 | A/starling/Rostov_on_Don/39/2007 |
| EPI_ISL_15358 | A/muscovy duck/Rostov_on_Don/51/2007 |
| EPI_ISL_63010 | A/chicken/Primorje/1/2008 |
| EPI_ISL_32668 | A/bean_goose/Tyva/10/2009 |
| EPI_ISL_32667 | A/grebe/Tyva/3/2009 |
| EPI_ISL_80265 | A/great_crested_grebe/Tyva/22/2010 |
| EPI_ISL_84604 | A/grebe/Tyva/2/2010 |
| EPI_ISL_31936 | A/black-headed gull/Tyva/115/2009 |
| EPI_ISL_28437 | A/chicken/Primorsky/85/2008 |
| EPI_ISL_9967 | A/chicken/Kurgan/3/2005 |
| EPI_ISL_64866 | A/duck/Novosibirsk/02/05 |
| EPI_ISL_10436 | A/duck/Tuva/01/2006 |
| EPI_ISL_10361 | A/chicken/Krasnodar/01/2006 |
| EPI_ISL_10360 | A/chicken/Mahachkala/05/2006 |
| EPI_ISL_64343 | A/common gull/Chany/P/2006 |
| EPI_ISL_31937 | A/great crested grebe/Tyva/120/2009 |
| EPI_ISL_25228 | A/falcon/Saudi_Arabia/D1936/2007 |
| EPI_ISL_15688 | A/Mergus albellus/Slovakia/Vh212/2006 |
| EPI_ISL_27786 | A/Peregrine falcon/Slovakia/Vh242/2006 |
| EPI_ISL_6414 | A/swan/Slovenia/760/2006 |
| EPI_ISL_27791 | A/cygnus olor/Slovenia/156/06 |
| EPI_ISL_27792 | A/Ardea cinerea/Slovenia/185/06 |
| EPI_ISL_27793 | A/Anas platyrhynchos/Slovenia/359/2006 |
| EPI_ISL_27794 | A/Anas acuta/Slovenia/470/06 |
| EPI_ISL_6249 | A/chicken/Sudan/1784_7/2006 |
| EPI_ISL_7014 | A/chicken/Sudan/2115_10/2006 |
| EPI_ISL_13230 | A/tufted duck/Sweden/V526/2006 |
| EPI_ISL_13231 | A/goosander/Sweden/V539/2006 |
| EPI_ISL_13233 | A/tufted duck/Sweden/V599/2006 |
| EPI_ISL_24705 | A/eagle owl/Sweden/V618/2006 |
| EPI_ISL_13240 | A/smew/Sweden/V820/2006 |
| EPI_ISL_24704 | A/eagle owl/Sweden/V1218/2006 |
| EPI_ISL_13249 | A/herring gull/Sweden/V1116/2006 |
| EPI_ISL_13247 | A/tufted duck/Sweden/V1027/2006 |
| EPI_ISL_13246 | A/tufted duck/Sweden/V998/2006 |
| EPI_ISL_13245 | A/canada goose/Sweden/V978/2006 |
| EPI_ISL_13241 | A/mute swan/Sweden/V827/2006 |
| EPI_ISL_69212 | A/common pochard/Switzerland/WV4080110/2008 |
| EPI_ISL_11231 | A/common coot/Switzerland/V544/2006 |
| EPI_ISL_11917 | A/tufted duck/Switzerland/V504/2006 |
| EPI_ISL_12822 | A/mute swan/Switzerland/V68/2006 |
| EPI_ISL_12823 | A/little grebe/Switzerland/V330/2006 |
| EPI_ISL_12824 | A/duck/Switzerland/V389/2006 |
| EPI_ISL_12825 | A/duck/Switzerland/V426/2006 |
| EPI_ISL_12826 | A/duck/Switzerland/V487/2006 |
| EPI_ISL_12827 | A/common pochard/Switzerland/V505/2006 |
| EPI_ISL_12828 | A/mallard/Switzerland/V537/2006 |
| EPI_ISL_12829 | A/common pochard/Switzerland/V558/2006 |
| EPI_ISL_12830 | A/common pochard/Switzerland/V592/2006 |
| EPI_ISL_12831 | A/common pochard/Switzerland/V762/2006 |
| EPI_ISL_10107 | A/turkey/Turkey/1/2005 |
| EPI_ISL_83693 | A/Pigeon/Turkey-Agri/09rs2841-8/2005 |
| EPI_ISL_83692 | A/Pigeon/Turkey-Kars/09rs2841-7/2005 |
| EPI_ISL_83700 | A/Chicken/Turkey-Edirne/09rs2843-5/2008 |
| EPI_ISL_83699 | A/Chicken/Turkey-Batman/09rs2842-92/2007 |
| EPI_ISL_83698 | A/Chicken/Turkey-Bitlis/09rs2841-93/2006 |
| EPI_ISL_83697 | A/Chicken/Turkey-Batman/09rs2841-71/2006 |
| EPI_ISL_83696 | A/Chicken/Turkey-Mus/09rs2841-51/2006 |
| EPI_ISL_83695 | A/Chicken/Turkey-Mus/09rs2841-50/2006 |
| EPI_ISL_83694 | A/Chicken/Turkey-Isparta/09rs2841-37/2006 |
| EPI_ISL_10340 | A/chicken/Crimea/08/2005 |
| EPI_ISL_103335 | A/swan/England/AV26-70/2008 |
| EPI_ISL_103350 | A/swan/England/AV380-2326/2008 |
| EPI_ISL_99850 | A/Common Buzzard/Berlin/1/2006 |
| EPI_ISL_6924 | A/chicken/Egypt/2253_1/2006 |
| EPI_ISL_6925 | A/turkey/Egypt/2253_2/2006 |
| EPI_ISL_6384 | A/duck/Ivory Coast/1787_18/2006 |
| EPI_ISL_2932 | A/chicken/Nigeria/SO494/2006 |
| EPI_ISL_66863 | A/chicken/Nigeria/08RS848-117/2007 |
| EPI_ISL_66864 | A/chicken/Nigeria/08RS848-118/2007 |
| EPI_ISL_66867 | A/chicken/Nigeria/08RS848-121/2007 |
| EPI_ISL_292 | A/whooper swan/Mongolia/3/05 |
| EPI_ISL_293 | A/whooper swan/Mongolia/4/05 |
| EPI_ISL_372 | A/whooper swan/Mongolia/2/06 |
| EPI_ISL_548 | A/common goldeneye/Mongolia/12/2006 |
| EPI_ISL_9940 | A/migratory duck/Jiangxi/2300/2005 |
| EPI_ISL_9939 | A/migratory duck/Jiangxi/2295/2005 |
| EPI_ISL_9938 | A/migratory duck/Jiangxi/2136/2005 |
| EPI_ISL_9937 | A/migratory duck/Jiangxi/1701/2005 |
| EPI_ISL_9936 | A/migratory duck/Jiangxi/1657/2005 |
| EPI_ISL_9935 | A/migratory duck/Jiangxi/1653/2005 |
| EPI_ISL_3903 | A/duck/Zhejiang/11/2000 |
| EPI_ISL_3900 | A/duck/Shanghai/35/2002 |
| EPI_ISL_3891 | A/duck/Fujian/19/2000 |
| EPI_ISL_68660 | A/chicken/Hong Kong/786-2/1997 |
| EPI_ISL_966 | A/Chicken/Hong Kong/728/97 |
| EPI_ISL_62720 | A/bar-headed goose/Mongolia/X54/2009 |
| EPI_ISL_62738 | A/whooper swan/Mongolia/8/2009 |
| EPI_ISL_76273 | A/whooper swan/Mongolia/1/2010 |
| EPI_ISL_76635 | A/whooper swan/Mongolia/21/2010 |
| EPI_ISL_93807 | A/duck/Hokkaido/WZ101/2010 |
| EPI_ISL_93808 | A/duck/Hokkaido/WZ83/2010 |
| EPI_ISL_131214 | A/chicken/Shimane/1/2010 |
| EPI_ISL_131215 | A/mute swan/Toyama/1/2010 |
| EPI_ISL_123277 | A/chicken/Pakistan-Khi/NARC-520/2008 |
| EPI_ISL_123318 | A/crow/Pakistan-Khi/NARC-11672/2008 |

**Table S3.** Summary profile of H5N1 HPAIV sequences isolated from reported outbreaks in Europe and the Russia between May 2005 and June 2010 (*N* = 277).

| Country | N.s<br>eq | Collection date | Host type |
| --- | --- | --- | --- |
| Austria | 20 | February 2006-April 2006 | Wild |
| Belgium | 1 | November 2008 | Wild |
| Bosnia | 1 | February 2006 | Wild |
| Bulgaria | 1 | March 2010 | Wild |
| Croatia | 1 | October 2005 | Wild |
| Czech Republic | 6 | March 2006-July 2007 | Wild & domestic |
| Denmark | 8 | March 2006-April 2006 | Wild |
| France | 2 | February 2006-June 2007 | Wild |
| Germany | 57 | November 2005-October 2008 | Wild, domestic, & Other |
| Hungary | 5 | February 2006-January 2007 | Wild |
| Italy | 3 | February 2006 | Wild |
| Poland | 9 | March 2006-December 2007 | Wild & domestic |
| Romania | 78 | October 2005-March 2010 | Wild, domestic, & Other |
| Russia | 51 | July 2005-January 2010 | Wild & domestic |
| Slovakia | 2 | February 2006 | Wild |
| Slovenia | 5 | February 2006-March 2006 | Wild |
| Sweden | 11 | March 2006-April 2006 | Wild |
| Switzerland | 13 | February 2006-February 2008 | Wild |
| Ukraine | 1 | May 2005 | Domestic |
| United Kingdom | 2 | January 2008 | Wild |

**Table S4.**
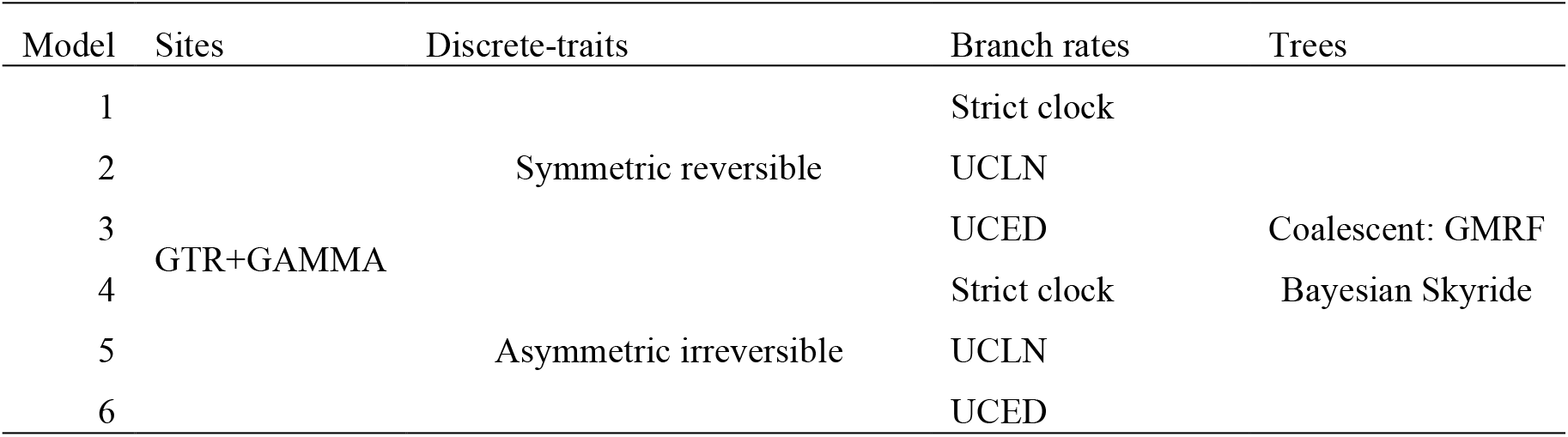
Candidate phylogeographic models for HA and NA gene segments explored for relative fit.

**Figure S1.**
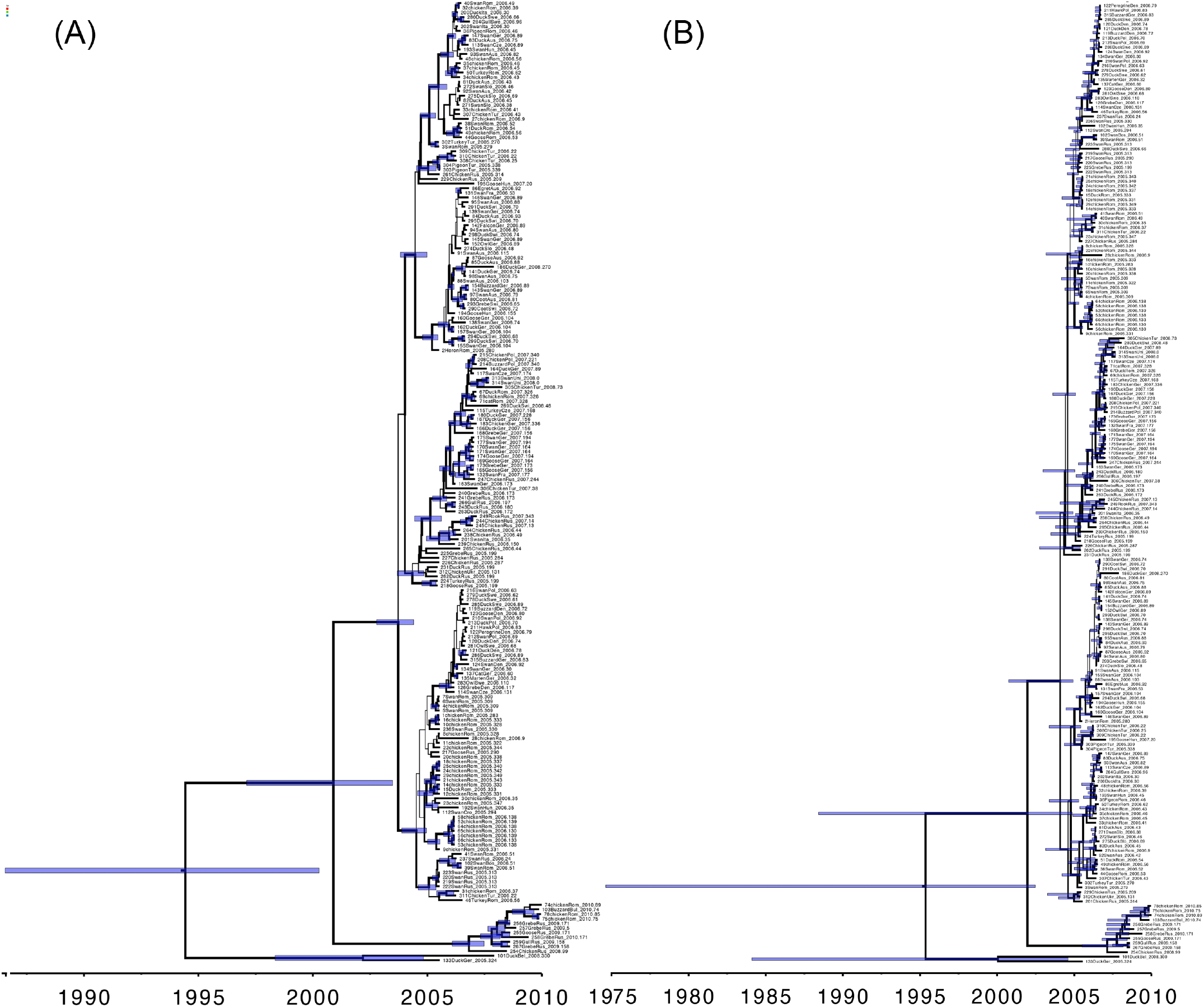
Maximum clade credibility (MCC) trees for H5N1 HPAIV hemagglutinin (HA) and neauraminidase (NA) gene regions (A and B, respectively). Branch lengths are rendered proportional to absolute time (see timescales), and the branch thickness is proportional to the posterior probability of the inferred host state. Nodes correspond to median ages and the blue horizontal bars at nodes represent the corresponding 95% HPDs for divergence-time estimates.

**Figure S2.**
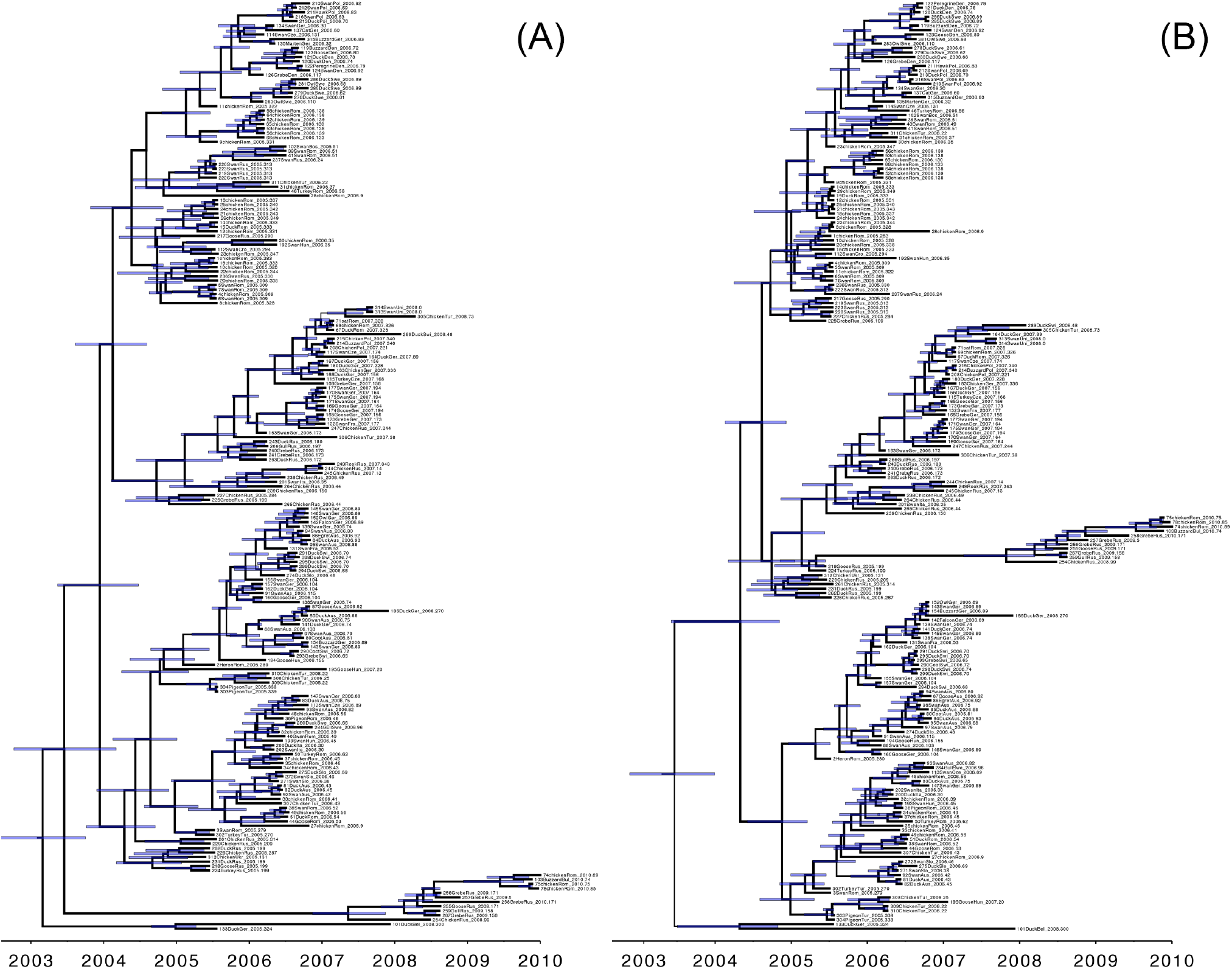
Maximum clade credibility (MCC) trees for H5N1 HPAIV hemagglutinin (HA) and neauraminidase (NA) gene regions (A and B, respectively). Branch lengths are rendered proportional to absolute time (see timescales), and the branch thickness is proportional to the posterior probability of the inferred location state. Nodes correspond to median ages and the blue horizontal bars at nodes represent the corresponding 95% HPDs for divergence-time estimates.

**Figure S3.**
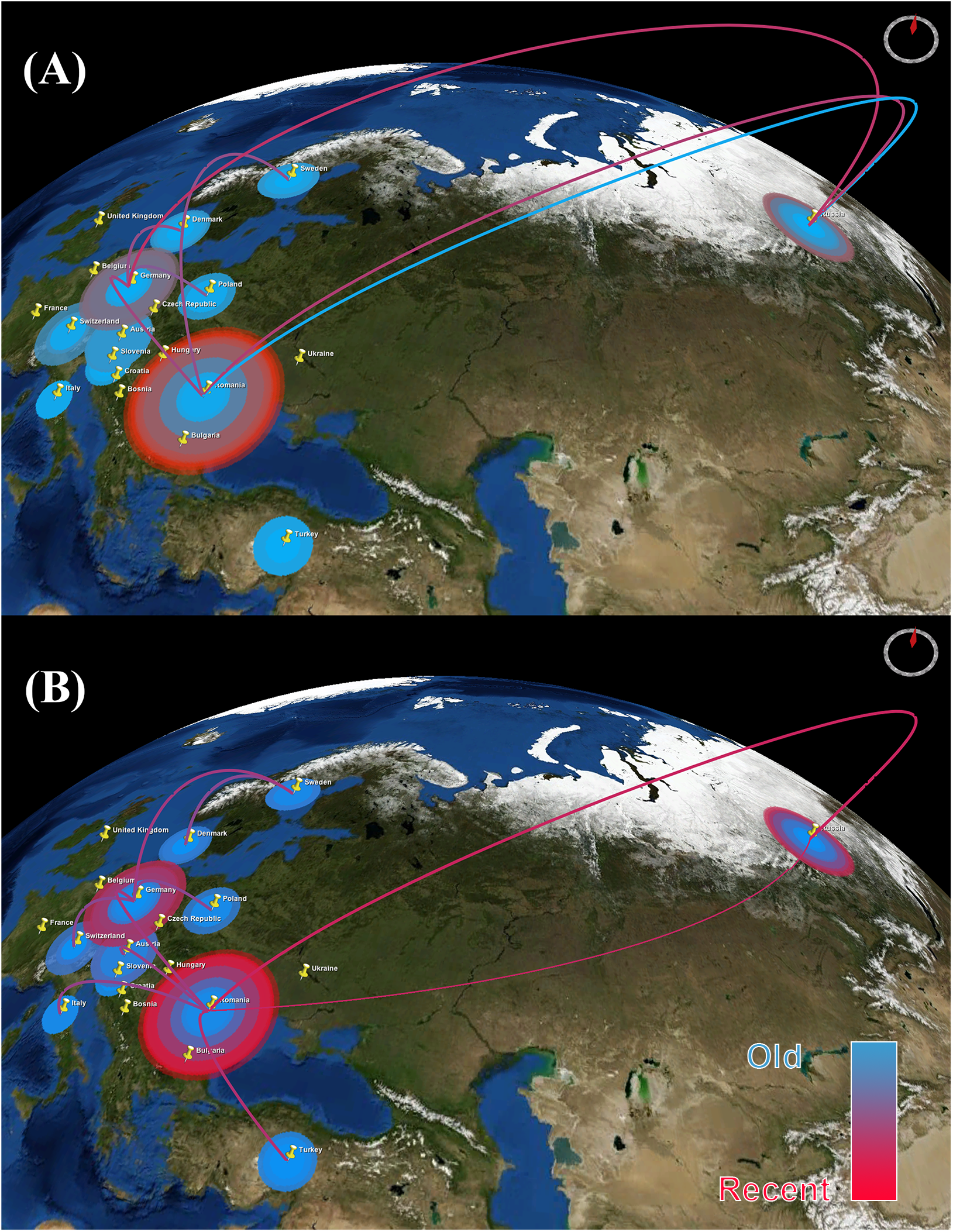
Temporal dynamics of the spatial diffusion of H5N1 HPAIV. The figure consists of snap shots of the dispersal pattern of avian influenza between May 2005 and June 2010. Lines between locations represent branches in the MCC tree along which spatial transmission occurs. The diameters of circles are proportional to square root of the number of MCC branches maintaining a particular location state at each time-point. The blue and red color gradients reflect the relative age of the transitions for each gene (older-recent, respectively). A: HA gene. B: NA gene. The maps are based on satellite pictures available in NASA World Wind (http://worldwind.arc.nasa.gov/java/).

**File S1 Temporal dynamics of the spatial diffusion of H5N1 HPAIV based on the HA gene.** The KML file demonstrates the dispersal pattern of avian influenza between May 2005 and June 2010. Lines between locations represent branches in the MCC tree along which spatial transmission occurs. The diameters of circles are proportional to square root of the number of MCC branches maintaining a particular location state at each time-point. The blue and red color gradients reflect the relative age of the transitions for HA gene (older-recent, respectively).

